# In vivo Biomedical Imaging of Immune Tolerant, Radiopaque Nanoparticle-Embedded Polymeric Device Degradation

**DOI:** 10.1101/2023.10.26.564238

**Authors:** Kendell M. Pawelec, Jeremy M.L. Hix, Arianna Troia, Matti Kiupel, Erik Shapiro

## Abstract

Biomedical implants remain an important clinical tool for restoring patient mobility and quality of life after trauma. While polymers are often used for devices, their degradation profile remains difficult to determine post-implantation. CT monitoring could be a powerful tool for in situ monitoring of devices, but polymers require the introduction of radiopaque contrast agents, like nanoparticles, to be distinguishable from native tissue. As device function is mediated by the immune system, use of radiopaque nanoparticles for serial monitoring therefore requires a minimal impact on inflammatory response. Radiopaque polymer composites were produced by incorporating 0-20wt% TaO_x_ nanoparticles into synthetic polymers: polycaprolactone (PCL) and poly(lactide-co-glycolide) (PLGA). In vitro inflammatory response to TaO_x_ was determined by monitoring mouse bone marrow derived macrophages on composite films. Nanoparticle addition stimulated only a slight inflammatory reaction, namely increased TNFα secretion, mediated by changes to the polymer matrix properties. When devices (PLGA 50:50 + 20wt% TaO_x_) were implanted subcutaneously in a mouse model of chronic inflammation, no changes to device degradation were noted although macrophage number was increased over 12 weeks. Serial CT monitoring of devices post-implantation provided a detailed timeline of device structural collapse, with no burst release of the nanoparticles from the implant. Changes to the device were not significantly altered with monitoring, nor was the immune system ablated when checked via blood cell count and histology. Thus, polymer devices incorporating radiopaque TaO_x_ NPs can be used for in situ CT monitoring, and can be readily combined with multiple medical imaging techniques, for a truly dynamic view biomaterials interaction with tissues throughout regeneration, paving the way for a more structured approach to biomedical device design.

## 1 Introduction

Biomedical devices fulfill many important roles in regenerative medicine: stabilizing injured tissue, supporting repair after trauma and restoring biological function. The materials chosen for implanted devices must be biocompatible and have the mechanical strength and stability to perform the intended task immediately post-implantation. An important aspect of device design is not only function at early time points, but compensating for the evolution of device properties over time. This is particularly relevant for devices that degrade in the body, producing degradation products that may also have physiological activity [1].

Even for well-characterized biocompatible materials, the degradation rate post-implantation can be variable due to the physiological environment [2,3]. Once implanted, device interactions are mediated by immune cells such as neutrophils and macrophages [4]. After the initial acute inflammatory reaction, macrophages are responsible for the majority of the foreign body response, and can promote revascularization and support fibrous encapsulation [5]. In addition, macrophages have been proposed to accelerate degradation of polymeric devices, as chronic inflammation has been linked to acidosis, for example within tumor microenvironments [6]. Extrapolating in vivo biomaterials degradation from in vitro degradation studies is problematic. To date, the majority of biomaterial degradation studies are conducted in buffers simulating a neutral physiological environment (saline, pH 7.4, 37°C), and in vivo degradation rates are significantly faster [7].

With in vivo material degradation remaining a black box, we have sought to utilize non-invasive longitudinal monitoring to understand the kinetics of material mass loss. Specifically, we have introduced radiopacity to several FDA-approved polymers by incorporating biocompatible tantalum oxide (TaO_x_) nanoparticle contrast agents into the polymer matrix [8-10]. This technique has been used to serially image porous tissue engineered devices via computed tomography (CT), a clinical imaging technique that can quickly and clearly distinguish implanted material from surrounding tissues [11].

To further validate that contrast agent incorporation does not significantly perturb biomaterials interactions post-implantation, we first investigated the in vitro response of macrophages to 0-20wt% TaO_x_ nanoparticle addition to synthetic polymers in use for medical devices: polycaprolactone (PCL) [12] and poly(lactide-co-glycolide) (PLGA) [13]. With nanoparticles stimulating only a low inflammatory response, radiopaque devices were implanted subcutaneously in mice, with and without chronic inflammation induced at the site, to determine if an in vivo effect of inflammation on implant degradation could be quantified. While serial monitoring is possible, contrast agent addition and the radiation exposure from CT can also influence the physiological environment around model implanted biomedical devices (phantoms), skewing reported degradation kinetics. For this reason, we have demonstrated that serial radiation exposure did not influence the time course of degradation. Together this supports the use of TaO_x_ NPs incorporated into polymers as a way to introduce imaging functionality into implanted medical devices, and potentially avoiding catastrophic device failure in the clinic through serial monitoring.

## 2 Materials & Methods

The study utilized three types of biocompatible polymers: polycaprolactone (PCL), poly(lactide-co-glycolide) (PLGA) 50:50 and PLGA 85:15. PCL (Sigma Aldrich) had a molecular weight average of 80 kDa. PLGA 50:50 (Lactel/Evonik B6010-4) and PLGA 85:15 (Expansorb® DLG 85-7E, Merck) were both ester terminated and had a weight average molecular weight of between 80-90 kDa.

Hydrophobic TaO_x_ nanoparticles were manufactured based on our previous procedure with one minor change [8]. Hydrophobicity is imparted by coating nanoparticles with hexadecyltriethoxysilane (HDTES, Gelest Inc., cat no SIH5922.0), an aliphatic organosilane. The coating has been shown to produce polymer composites with homogeneous nanoparticle distributions [11].

### 2.1 Phantom manufacture

Phantoms were created with micro-scale porosity (< 100μm) and with macro-scale porosity (200 - 500 μm) to mimic tissue engineering constructs which must accommodate both nutrient diffusion and cell and tissue infiltration [11]. For in vitro experiments, films mimicking the surface of the phantom struts was created. To determine if macrophage response to polymer composites was universal, three types of biocompatible polymer films were produced. Prior to casting films, polymers were solubilized in suspensions containing TaO_x_ nanoparticles in dichloromethane (DCM, Sigma); PCL was used at 4wt %, while PLGA solutions ranged from 10wt% (0-5wt% TaO_x_) to 8wt% (20wt% TaO_x_). The amount of TaO_x_ (0-20wt%) was calculated based on the weight percent of the total dry mass (polymer + nanoparticle). To introduce microporosity to the films, sucrose (mean particle size 31 ± 30 μm, Meijer) was added to the solution, so that the final slurry was 70 vol % sucrose and 30 vol % polymer+nanoparticles. Suspensions were vortexed, then cast and dried on glass sheets. Dried films were removed and submerged in Milli-Q water to remove the sucrose, then dried for storage. In addition, for cell culture, non-porous PCL films were created in the same way, without the addition of nanoparticles or sucrose.

For 3D phantoms, PLGA 50:50 was solubilized in a suspension of TaO_x_ nanoparticles in DCM to make a 12wt% PLGA solution. Sucrose was added to the suspension, calculated to be 70 vol% of the polymer + nanoparticle mass in solution, as before, followed by NaCl (Jade Scientific) at 60 vol% of the total polymer + nanoparticle volume. The suspension was vortexed for 10 minutes and pressed into a silicon mold that was 4.7 mm diameter, 2 mm high. After air drying, phantoms were removed, trimmed of excess polymer and then washed for 2 hours in distilled water, changing the water every 30 minutes to remove sucrose and NaCl. Washed phantoms were air dried overnight and stored in a desiccator prior to use.

### 2.2 Macrophage culture

#### Cell harvest

Bone marrow cells were isolated from the femurs of adult Balb/c mice (male and female), euthanized for other purposes [14]. The femurs were placed in 70% ethanol and then kept on sterile ice-cold PBS. The ephiphyses of each bone was removed with scissors and the bone marrow was flushed with 5 ml PBS+1% Penicillin–Streptomycin (ThermoFisher Scientific, 15,140,122), by inserting a 22G needle into the bone shaft and collecting the hydrate in a sterile centrifuge tube. The collected cells were centrifuged for 5 min at 350g and resuspended in 500 μl of PBS. Red blood cells were lysed with 0.8% ammonium chloride (Fisher Scientific, NC9041684) for 10 minutes on ice. The bone marrow cells were resuspended in PBS and passed through with a 70 μm cell strainer. The cells either used immediately, or resuspended in freezing media (90% fetal bovine serum (FBS, Life Technologies, 10438018) + 10% dimethyl sulfoxide (DMSO, Sigma)) and stored in liquid nitrogen until use. The bone marrow of 2-3 individuals was combined into each lot of cells.

Prior to starting in vitro experiments, bone marrow cells were differentiated into bone marrow derived macrophages. Bone marrow cells were transferred to non-tissue culture treated flasks and grown at 37∘C and 5% CO_2_ in differentiation media (RPMI 1640+GlutaMAX (Gibco, 61870036), 10% heat-inactivated, low endotoxin FBS (Life Technologies, A3840201) + 20 ng/ml recombinant mouse macrophage colony-stimulating factor (M-CSF, R&D systems, 416ML010)) for 6-7 days. The cells which were adherent after differentiation were considered macrophages.

#### Preparation of substrates

Disks of porous films were punched (13 mm diameter) and sterilized by immersion in 70% ethanol for 30 min, followed by two washes with sterile water. Sterile, hydrated films were placed into 24-well inserts (CellCrown™, Z681903-12 EA), as described previously [15]. Briefly, each sample consisted of two layers: a bottom layer of non-porous PCL (punched to 24mm diameter) and a top layer of porous film. After assembly into the inserts, they were placed at the bottom of a 24 well tissue culture plate, and the entire assembly was placed in UV light for 30 min.

#### In vitro culture

Differentiated macrophages were passaged with 0.5% Trypsin-EDTA (Gibco, 25300-054), and plated onto inserts at 2-3×10^4^ cells/well in 400 μl of differentiation media. Media was changed every 2-3 days and the supernatant was collected for testing at day 7 and 14 post-seeding. Protein was harvested at day 14. As controls for macrophage polarization, cells were also seeded on tissue culture treated polystyrene at the same density. Two days prior to harvesting supernatant or protein, control cells were treated with media supplemented with polarization factors, to serve as positive controls for assessing polarization on film surfaces. The three control groups were 1) no polarization (M0) (unsupplemented differentiation media), 2) pro-inflammatory M(LPS/IFNγ) (differentiation media supplemented with 100 ng/ml lipopolysaccharide (LPS, Sigma, L4391) and 50 ng/ml interferon gamma (IFNγ, Pepro Tech, 315-05-20UG)) or 3) anti-inflammatory M(IL4/IL13) (differentiation media supplemented with 40 ng/ml IL-4 (Pepro Tech, 214-14-5UG) and 20 ng/ml IL13 (Pepro Tech 210-13-2UG)). All tests were done in triplicate from different cell lots, with two technical replicates each, and presented as mean ± standard error.

### 2.3 Macrophage polarization

Macrophage polarization in response to culture on polymer composites was assessed in several ways. Supernatant was investigated at day 7 utilizing ELISA kits, following manufacturer’s instructions. Supernatant was probed for tumor necrosis factor alpha (TNFα, R&D Systems, mta006), IL10 (Invitrogen, BMS614INST), and CCL17 (R&D Systems, mcc170). On day 7, cellular levels of arginase I were also quantified by ELISA (abcam, ab269541), lysing cells in protein lysis buffer for 15 minutes on ice with the reagents provided by the kit, following manufacturer’s instructions. In addition, the nitrite concentration in the supernatant was measured via the Griess assay (Promega, TB229), per manufacturer’s instructions. In all cases, supernatant from control macrophages, polarized within 48 hours, were run alongside the film samples.

At day 14, sodium dodecyl sulfate polyacrylamide gel electrophoresis (SDS-PAGE) and western blotting was utilized to probe the expression of inflammation markers. Protein was harvested from macrophages by incubating films in Ripa buffer overnight, at 4°C while shaking. The protein concentration was measured (BioRad, BCA kit) and samples with 2× Laemmli buffer were incubated at 60°C for 20 min before separation by SDS/ PAGE. After transfer to PVDF membranes (iBlot 2 Gel Transfer Device, invitrogen), membranes were blocked with 5% dried milk in phosphate buffered saline (PBS) + 0.1% Tween-20 (PBST) and probed with primary antibodies specific for IRF5 (1:875, abcam, ab181553), TGFβ1 (1:700, abcam, ab215715), or integrin α_V_(1:1100-1:1400, abcam, ab302640) at 4 °C overnight. The next day, the membrane was washed and incubated with anti-mouse IgG-HRP (12-349, Millipore) or anti-rabbit IgG-HRP (ab205718, abcam) at room temperature for 2 hr. Protein bands were visualized with Amersham ECL Plus substrate (cytiva, RPN2232) on a Li-Cor Odyssey FC and resulting signal was quantified via Image J.

Developed membranes were washed in PBS and stripped for 15 minutes in Restore Western Blot Stripping Buffer (Thermo Scientific 21059) before being either blocked and reprobed or stained for total protein using the BLOT-Fast Stain (G Biosciences, cat# 786-34) according to manufacturer’s directions. The total protein signal was imaged under visible light on a c300 imager (azure biosystems). Protein expression was quantified via Image J and normalized to total protein to adjust the measured band intensity. For comparisons between membranes, all signals were normalized to PCL + 0wt% TaO_x_, which was run on every membrane. All data reported are the result of three independent replicates (two technical replicates each) and are reported as mean ± standard error. The full blots are presented in Supplemental data (Section S3).

### 2.4 Microscopy

Macrophages on films and glass coverslips were fixed in 4% paraformaldehyde and stored in PBS at 4 °C. For staining, films were washed in PBST, then permeabilized in 0.1% Triton X-100 in PBS, followed by two PBST washes. Samples were then blocked with 5 wt% bovine serum albumin (BSA) in PBST, for 1 hr, at room temperature. After washing twice in PBST, films were incubated with primary antibodies, overnight at 4°C, in 3 wt% BSA/PBST. Following primary incubation and two PBST washes, secondary antibodies in PBS were incubated with the samples for 1–2 h at room temperature. Samples were then incubated for 30 minutes in ActinRed^TM^ 555 ReadyProbes^TM^ (Invitrogen, R37112, 2 drops per ml PBS) to visualize the actin cytoskeleton. Finally, films were washed twice with PBS and left in a solution of PBS and DAPI stain (1:000 in PBS, ThermoFisher Scientific, 62248). Primary antibodies, and the dilutions used, were: IRF5 (1:500, abcam, 181553), CD68 (1:100, invitrogen, MA5-13324). Secondary antibodies: Alexa Fluor 488 (1:1000, A11070), Alexa Fluor 647 (1:1000. A21236).

Stained films were inverted onto a coverslip in a drop of Slow Fade Diamond Antifade Mountant (Invitrogen, S36972). Imaging was performed using a Leica DMi8 microscope, using an LASX software interface. With the high surface roughness of the films, z-stacks were taken at a minimum of 3 places on the film and post-processed with Thunder image analysis (Leica) to remove background fluorescence.

Scanning electron microscopy (SEM) images were taken of films after 1 week of culture with cells and after the cells had been removed. To prepare for SEM, cells were fixed with paraformaldehyde as before and then dried in serial ethanol dilutions, at least 1 hour incubation in each: 70% ethanol, 80% ethanol, 90% ethanol, 100% ethanol. Finally samples were incubated in hexamethyldi-silazane (Sigma) for 10 minutes and allowed to air dry. Dried samples were adhered to 13 mm aluminum stubs and sputter coated with platinum. Surfaces were examined using a Zeiss Auriga, at 2 keV in scanning electron mode.

### 2.5 Molecular Imaging

#### Micro-computed tomography (µCT)

All tomography images were obtained using a Perkin-Elmer Quantum GX. Prior to implantation, phantoms were imaged at 90 keV, 88 µA, with a 25 mm field of view at a 50 µm resolution. After acquisition, individual phantoms were sub-reconstructed using the Quantum GX software to 15 µm resolution. Hydrated phantoms used for serial monitoring were imaged 24 hrs before implantation.

In vivo μCT on mice was performed at 90 keV, 88 μA. At each time point, one scan was taken of the phantom. The scan was 25 mm field of view (4 min total scan time) at 50 μm resolution. During acquisition, mice were anesthetized using an inhalant anesthetic of 1–3% Isoflurane in 1 L min−1 oxygen. Mice (n = 12) were scanned pre surgery for baseline, immediately post-implantation, on day 1 post-implantation, and at weeks 1, 2, 4, 6, 8, 10, 12 post-implantation. Total cumulative radiation dosage was 25-27 Gy over 12 weeks and was delivered to local regions around the phantoms. Additionally, scans were performed immediately following euthanasia at week 6 (n = 6) and week 12 (n = 27). The physiological result of radiation due to localized scans is dependent on the sensitivity of surrounding tissues such as adipose tissue or bone marrow [16]. In the present study, this was tracked via complete blood count and clinical pathology following euthanasia (Table S1 and S2, Supplemental).

From the tomography scans of phantoms, several parameters were quantified. 12-18 µm sub-reconstructions were used for all phantom analysis. Analysis of the polymer matrix component of phantoms was performed using ITK SNAP [17]. The polymer matrix was segmented from the background tissue and the volume of the matrix and its average intensity was calculated. Gross features of the phantoms (thickness, diameter) were analyzed using Image J for scaffolds pre-implantation and at weeks 0-4, and ITK-SNAP was used to report gross volume for weeks 6-12, utilizing a different segmentation from that used for polymer matrix calculation. In Image J, image stacks were opened and rotated, using built in functions, to measure thickness and diameter in 5 planes, which were averaged. From the diameter and thickness, a “gross volume” was defined as the volume occupied by a solid cylinder with the corresponding thickness and diameter. Regardless of the program used for calculation, the gross volume of phantoms consisted of both the polymer matrix volume and the volume of the pores. From this, percent porosity of the phantoms was calculated as the percentage of the gross volume not occupied by matrix. All data quantified from μCT scans is reported as mean ± standard deviation.

#### Magnetic Resonance Imaging (MRI)

Mice were scanned on a 7T preclinical MRI (Bruker, Billerica, MA). Full body scans were acquired for mice using a 72 mm volume coil. 3D T1_FISP scan parameters were: TR/TE 6/3 ms, 120×70×64 mm FOV, 500 um resolution, 6 averages, 11 min scan. MRI scans were obtained at week 5 for comparison to μCT mid study.

### 2.6 In vivo inflammation study

#### Phantom preparation

Phantoms of PLGA 50:50 incorporating 20 wt% TaO_x_ nanoparticles were prepared as detailed above. After preparation, phantoms were cut to 3 mm diameter with a biopsy punch and were soaked in 70% ethanol for 30 min. The ethanol was replaced by sterile PBS and centrifuged for 10 minutes at 11,000 rpm. The PBS was replaced and the implants were left in sterile PBS at 37 °C until implantation. As a comparison to published literature on PLGA 50:50 degradation, a further set (n=3) of phantoms with 20wt% TaO_x_ were hydrated, as before, and placed in sodium citrate at pH 5.5 for 12 weeks. The degradation buffer was changed every week, and scaffold degradation was tracked via µCT imaging.

#### Surgical implantation

All procedures were performed in accordance with IACUC-approved protocols and Veterinary guidelines at Michigan State University. BALB/c Mice (n = 45 adult males, 2 months old; Charles River Laboratories) were used for this surgical implantation and μCT imaging study. Mice (n = 30) were surgically implanted with one PLGA 50:50 phantom containing 20 wt% TaO_x_ nanoparticles installed within a single implantation surgical site. The PLGA 50:50 phantoms were placed subcutaneously dorsal-thoracic between the scapulae. Mice (n = 12) were given a sham surgical procedure that consisted of placing a suture only without the implant. Mice (n = 3) were kept as naive controls. Each animal implanted or receiving a sham procedure was administered analgesia at least 20 min prior to making the initial incision, including prophylactic Ampicillin (25 mg kg−1; SID) administered S.C., Meloxicam (5 mg kg−1; SID) administered S.C. in the right dorsal lateral flank of the animal, and local infiltration of 2% Lidocaine (diluted to 0.5%) was administered S.C. along the intended incision site just prior to making the cutaneous incision (7 mg kg−1 Max dose; SID). Animals were anesthetized via inhalant isoflurane (3–4% isoflurane in 0.8–1 LPM oxygen for induction) and maintained via inhalant isoflurane during surgery (1–3% isoflurane in 0.8-1 LPM oxygen). Supplemental heat was provided via recirculating warm water blankets during anesthesia induction, patient preparation, surgery, and patient recovery. Each implant was sutured to surrounding tissue with at least one single interrupted suture using 9-0 PROLENE (Polypropylene) monofilament Suture for in situ location retention. Incision sites were closed in multiple layers where necessary using 5-0 COATED VICRYL (polyglactin 910) Suture and cutaneous layers were closed using 5-0 PDS-II (polydioxanone) Suture. Following animal recovery, Meloxicam (5 mg kg−1; SID) and Buprenorphine (2 mg kg−1, BID, every 8–12 h) were administered S.C. in the left or right dorsal lateral flank of the animal for 48 h following surgery. Post-operative clinical observation, body weight assessment, and health score assessment were performed for 7–14 days postoperatively.

#### Carrageenan preparation and administration

Carrageenan (Sigma Aldrich, 22049) was formulated to a 1.7wt% solution and autoclaved in a sterile glass vial with rubber septum, by dissolving 187 mg Carrageenan in 11 mL diluent (sterile saline, 0.9% Sodium Chloride for Injection, USP) at 80°C. The formulated carrageenan solution was administered through subcutaneous (SC) injection by tenting the animal’s skin in the dorsal thoracic region between the scapulae at the surgical implantation site and inserting a 25G 5/8"-27G 1/2" needle through the epidermis and into the SC "pocket" of the tent. A single bolus dose was administered at week 3 and again at week 4 post-surgical implantation. Animals (n = 21) dosed with formulated carrageenan at these two time points exhibited clinical signs of inflammation distal to the surgical implantation site, therefore additional carrageenan dosing was discontinued and animals were allowed to recover for the remainder of the study.

#### End of study

Euthanasia was performed on week 6 (n = 6) and week 12 (n = 39). At termination, animals were sacrificed via CO_2_ overdose asphyxiation. Following euthanasia, blood samples were collected for complete blood count and chemistry panel (Table S1-2, Supplemental Information). Tissue samples of the heart, brain, kidney, bladder, liver, and spleen were also collected post-euthanasia for histopathological hematoxylin and eosin (H&E) staining to investigate for signs of inflammation.

**Figure 1:**
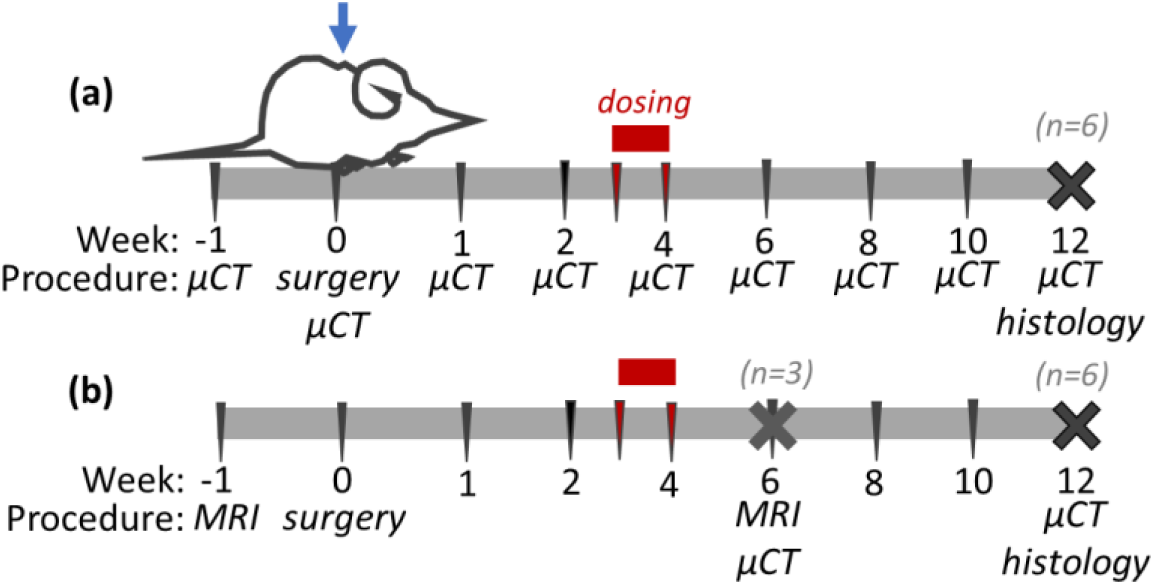
Degradation of radiopaque phantoms, mimicking biomedical implants, was tracked over 12 weeks after subcutaneous implantation. To induce chronic inflammation, groups were dosed over weeks 3-4 with 1.7wt% carrageenan or 0wt% carrageenan (control). (a) Serial monitoring of implants via CT was compared to (b) controls imaged only postmortem at discrete time points; at all points where CT was performed for serially monitored groups, control groups underwent sham imaging to ensure no effects of animal handling.

### 2.7 Statistics

Statistics were performed using GraphPad 9.4.1. Data was analyzed via ANOVA, followed by Fishers LSD test. In all cases, α < 0.05 was considered significant, with a 95% confidence interval.

## 3 Results

Macrophages are expected to be the key immune cell type to interact with devices post-implantation. Accordingly, macrophage response was probed after culture on films of biocompatible polymers incorporating nanoparticles. BMDM macrophages could be polarized towards a pro-inflammatory and pro-healing phenotype, M(LPS/IFNγ) and M(IL4/IL13), respectively, expressing markers associated with each [18]. M(LPS/IFNγ) macrophages had a high significantly higher concentration of TNFα and nitrite in the supernatant. M(IL4/IL13) polarized cells had high levels of CCL-17 and Arginase-1 and expressed significantly more TGFβ1 (Supplemental S1). Expression of IL10 was not significantly altered during polarization.

### 3.1 Immune response to nanoparticle composites

In all cases, macrophages were able to adhere to the porous film surfaces and the adhesion integrin α_V_ was differentially expressed across film types and with different amount of nanoparticle incorporation, Figure 3(c). Expression of integrin α_V_ decreased with nanoparticle addition, even at 5wt% TaO_x_ on PLGA films. After one week of culture, the surface of PLGA 50:50 films, the fastest degrading of the polymers tested, had pitting regardless of nanoparticle incorporation, Figure 2(e-f). The macrophages in culture were responsible for the change in surface morphology, as incubation in media alone at 37°C had no affect on the surface. No other films experienced the same change in surface morphology (Supplemental Figure S2).

**Figure 2:**
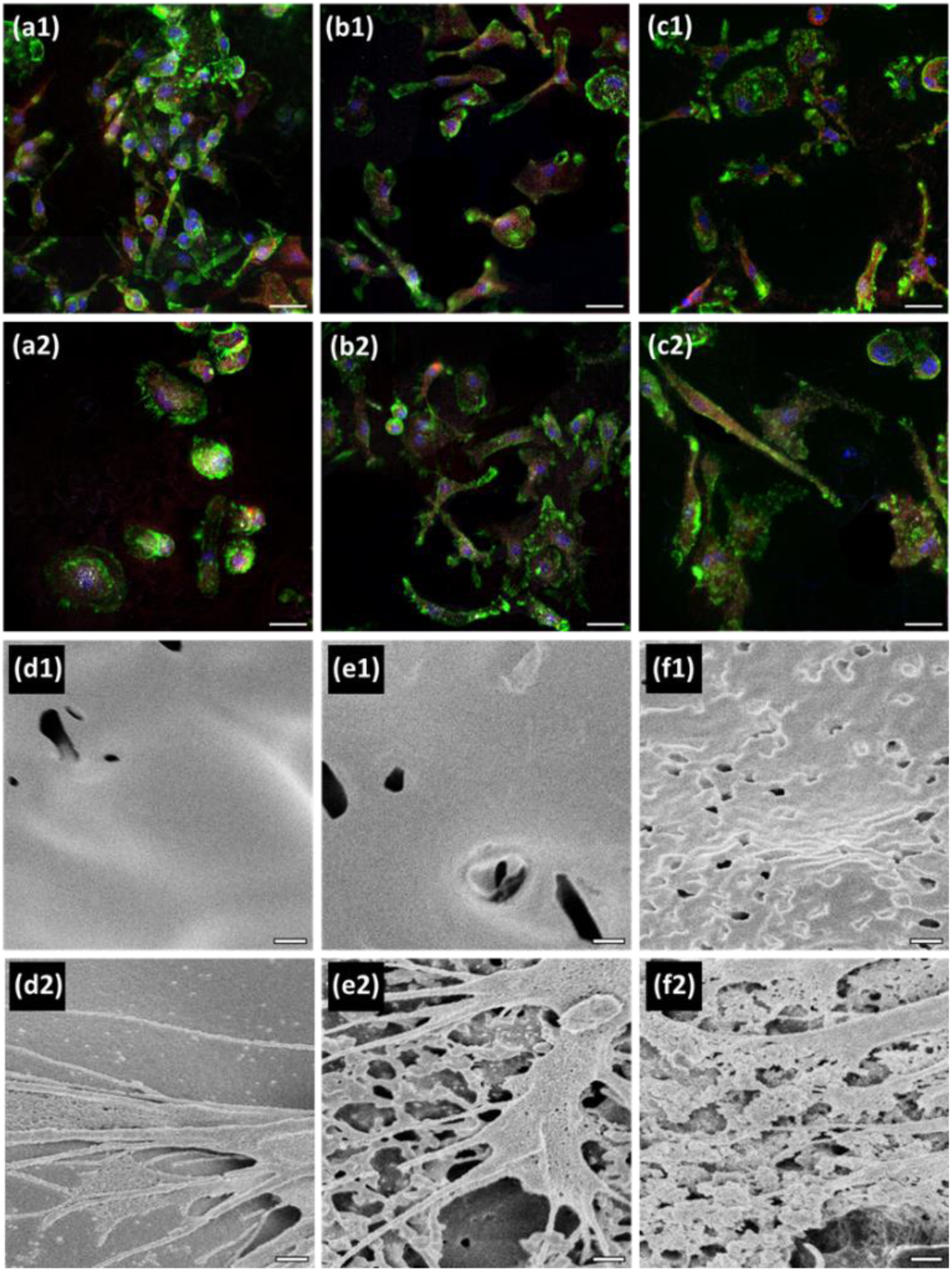
Bone marrow derived macrophages attached and proliferated on all film surfaces: (a) PCL, (b) PLGA 85:15 and (c) PLGA 50:50 with both (1) 0wt% TaO_x_ and (2) 20wt% TaO_x_ incorporation. Green: actin cytoskeleton, red: CD68, white: IRF5, blue: nucleus (DAPI). Film surface morphology was influenced by macrophage culture for fast degrading polymers, but not for others, as seen via scanning electron microscopy: (d) PLGA 85:15 + 0wt% TaO_x_, (e) PLGA 50:50 + 0wt% TaO_x_ and (f) PLGA 50:50 + 20wt% TaO_x_, imaged after one week of culture (1) without macrophages and (2) with macrophages seeded at the surface. Scale bar: (a-c) 25 μm, (d-f) 400 nm.

**Figure 3:**
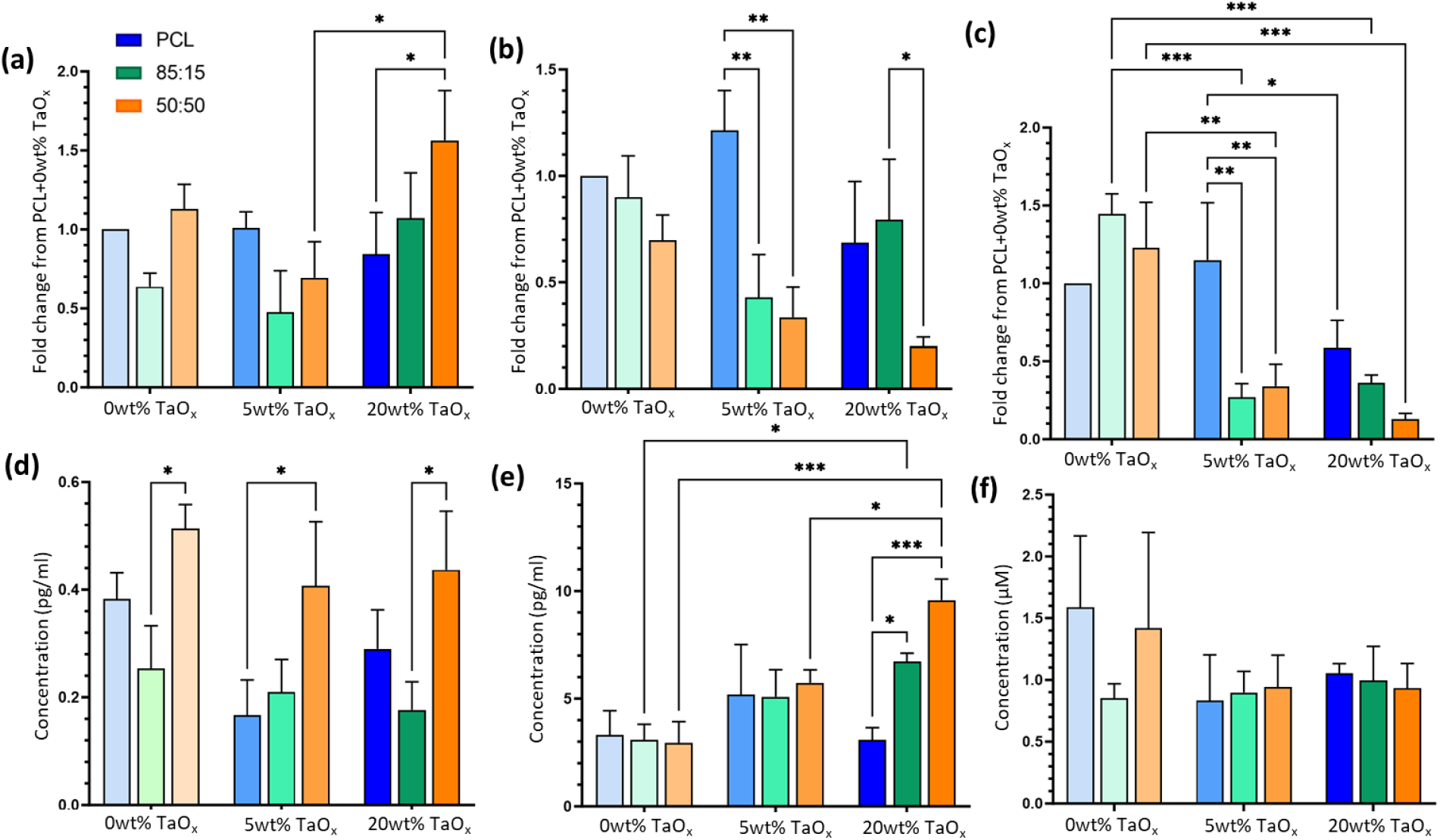
Macrophage polarization and attachment markers were significantly affected by polymer matrix and TaO_x_ nanoparticle content: (a) TGFβ1, (b) IRF5, (c) integrin α_V_, (d) CCL-17, (e) TNFα and (f) Nitrite. Data shown as mean ± standard error. *p < 0.05, **p < 0.01, ***p < 0.001.

Many of the changes in supernatant release and protein expression of BMDMs were tied to the polymer matrix rather than the addition of nanoparticles. However, a clear trend in cytokine release with nanoparticle addition was observed with secreted TNFα, a pro-inflammatory marker. On PLGA films, TNFα concentration increased as the amount of nanoparticles increased, Figure 3(a). Macrophages on PCL had the highest expression of IRF5, regardless of nanoparticle incorporation. In general, cytokines, such as CCL-17, and nitrite were secreted into the media only at very low levels, compared to polarized control macrophages.

### 3.2 Effect of inflammation on implant degradation

The systemic effects of inflammation on implant degradation were tracked by implanting phantoms subcutaneously in mice and observing the response after 6 and 12 weeks. The inflammatory response was compared to controls which received sham implants (surgery and suture at the implant site only). To mimic chronic inflammation and assess the impact of a longer-term immune response on implant degradation, an intense inflammation response was induced with injection of carrageenan at the implant site. When mice underwent sham surgery, the carrageenan successfully triggered a prolonged foreign body response, seen by increased granulomatous cellulitis around the implant site, Figure 4(f1-f2). The administration of carrageenan also caused inflammation and lesions at areas adjacent to the injection site.

**Figure 4:**
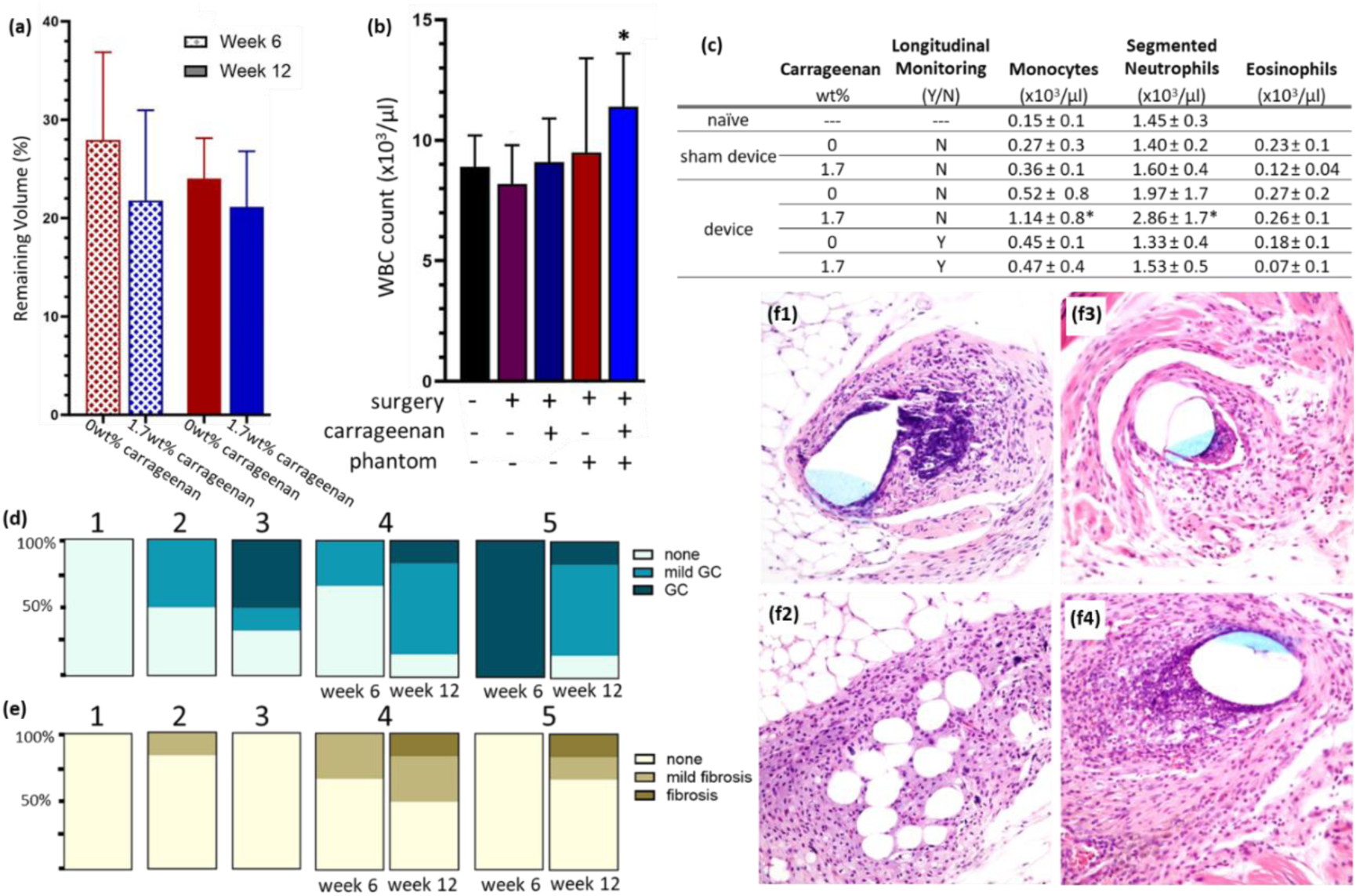
A chronic inflammation response could be induced in a mouse model, which was further intensified with the implantation of a phantom device. (a) Over 12 weeks the volume of the phantom matrix remaining (quantified from post-mortem CT scans) decreased. (b) Systemic changes in white blood cell (WBC) count were observed and (c) further quantified by monocytes, neutrophils and eosinophils. (d) Inflammation in the form of granulomatous cellulitis (GC) was noted at the surgical site, and with increased intensity around implanted phantoms. (e) Fibrosis was also present around implanted phantoms. The sample groups were 1: naive control, 2: no device, 0wt% carrageenan, no CT (week 12), 3: no device, 1.7wt% carrageenan, no CT (week 12), 4: device 0wt% carrageenan, no CT (weeks 6 and 12), 5: device, 1.7wt% carrageenan, no CT (weeks 6 and 12). (f) Representative histology (H&E staining) at the implantation site at week 12: 1: no device, 0wt% carrageenan, no CT, 2: no device, 1wt% carrageenan, no CT, 3: device 0wt% carrageenan, no CT, and 4: device, 1.7wt% carrageenan, no CT. * significantly different from all other groups (p < 0.05) Phantoms experienced degradation over the 12 weeks of implantation. By week 6 any distinct porosity was lost in the phantom and collapse continued through the remaining weeks, until only 10% of the pre-implanted gross volume of the phantom remained. This corresponded to a remaining matrix volume of 24 ± 3.7% and 21 ± 5.1%, with and without an additional immune stimulation, respectively (Figure 4(a)). While the change in matrix volume between weeks 6 and 12 was not significant, this is likely due to the smaller group size at week 6 (n=3) required to support end-point analysis techniques. Stimulating chronic inflammation did not significantly alter the timeline of matrix release from the implant site.

To simulate the worst case scenario of a biomaterial induced inflammatory response, the biocompatible polymer with the fastest degradation time and which was visibly acted upon by macrophages in vitro (PLGA 50:50) was chosen as the phantom, with 20wt% TaO_x_ incorporated. Implantation of a device provoked a larger local immune response compared to sham controls, with increased granulomatous inflammation, most apparent just after the carrageenan dosing at week 6, Figure 4(d). By week 12, animals that had an implanted phantom had similar inflammatory responses, regardless of carrageenan dosing. In the later stages of the foreign body response, related to extra-cellular matrix (ECM) deposition, or fibrosis, implantation of a device increased scarring. Even by 6 weeks post-implantation, only the suture was clearly distinguishable from tissue at the implant site.

Over and above the inflammatory response at the local implant site, placement of the devices had a systemic influence, tending to increase the total circulating white blood cell (WBC) count at 12 weeks, Figure 4(b). Without a device present, the response to carrageenan was a mild increase in circulating white blood cells, which was significantly increased with a device present. The increase in white blood cells was tied to a significant increase in the number of monocytes, the cells that replace tissue specific macrophage populations, and neutrophils, Figure 4(c). Additionally, with surgery and device implantation, a measurable number of eosinophils was present, not observed in naive control animals. Having TaO_x_ present in the implanted phantoms did not induce any pathological damage to target organs or excretory organs, Supplemental S4.

### 3.3 Tracking device degradation in situ

With 20wt% TaO_x_ incorporation, phantoms were visible via CT throughout implantation, indicating that the nanoparticles did not disperse from the structure prior to collapse, Figure 5. Also, there was no observation of any widespread clouding of the tissue surrounding the devices that might be indicative of a burst release of the nanoparticles from the polymer matrix. As demonstrated previously [10], the addition of radiopaque contrast agents does not create artifacts in other medical imaging techniques, such as MRI, Figure 5(f).

**Figure 5:**
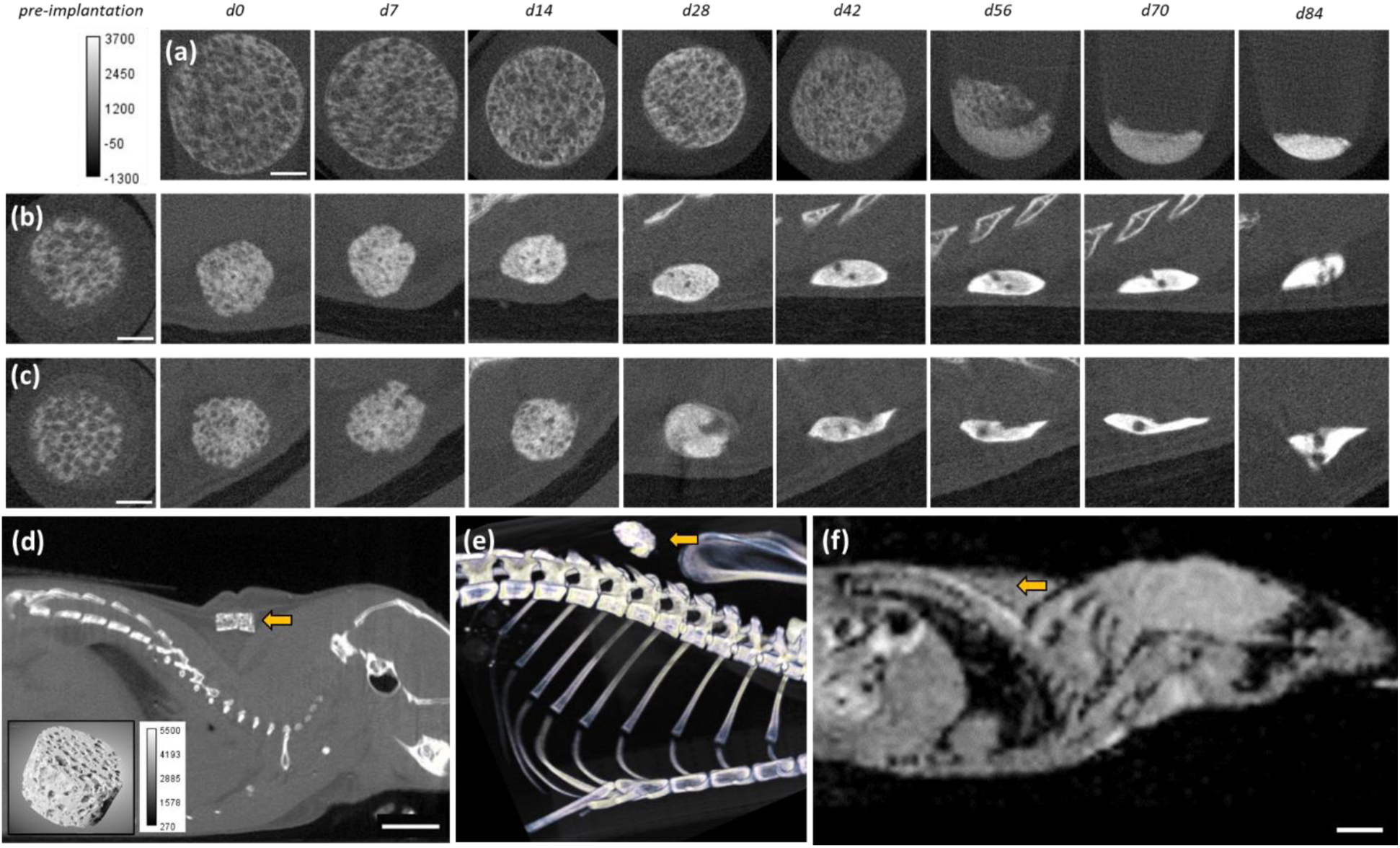
Addition of radiopaque contrast agents into polymer matrices allowed for the serial monitoring of phantoms post-implantation. Phantoms of PLGA 50:50 + 20wt% TaO_x_ first lost their porous structure and then compacted. The timeline was affected by the environment: (a) in vitro degradation at pH 5.5 (b) in vivo degradation with 0% carrageenan and (c) in vivo degradation with 1.7wt% carrageenan. (d) The phantoms are clearly visible from surrounding tissue, as seen via (d) CT scanning (inset: 3D rendering of the phantom) and (e) a 3D rendering. (f) While visible in CT, the phantoms were not discernible in MRI. Yellow arrow marks the implanted phantom. Scale bar (a-c): 1mm, (d-f): 5 mm.

The ability to serially monitor scaffolds over time allowed for a more complete picture of the device collapse and degradation over 12 weeks. Implanted phantoms in vivo completely lost their structure between days 28-42. This was significantly earlier than corresponding phantoms in vitro, even when the degradation test was conducted under conditions mimicking a harsh the lysosome environment (pH 5.5), Figure 5(a). Degradation responses in vivo are expected to be affected by cellular responses such as protease release, mechanics at the site, and fluid flow.

Longitudinal monitoring of the phantom collapse showed compaction of the phantom matrix and a loss of the distinct porous features engineered into the phantom to mimic tissue engineering devices. During the degradation of the phantoms, the compaction of the matrix increased the apparent radiopacity, by physically increasing the nanoparticle concentration, Figure 6(c). The greatest change in matrix volume was a consequence of implantation, likely due to the constraint of the phantom by surrounding tissues. If tracking only changes to the matrix that occurred in vivo (post-implantation), the phantoms retained 40-41% matrix volume over 12 weeks, rather than an apparent 21-24% if normalizing to the pre-implantation volume. However, to compare phantoms across groups, the pre-implantation volume was used for all reporting. Serially monitoring phantom degradation via CT did not affect the amount of remaining matrix in the phantom, with and without carrageenan. At week 12, the volume remaining with CT monitoring was 21 ± 1.2% and 23 ± 3.3% (with and without chronic inflammation), within the 21-24% range of phantoms that were no subjected to serial CT scans, Figure 6(a). This clearly demonstrated that monitoring can be achieved without altering the physiological processes involved in degradation. Serial monitoring also allowed tracking of implant porosity over time (Figure 6(b)), until phantoms reached 20-25% porosity, at which point the distinct porous structure disappeared, Figure 5(b-c).

**Figure 6:**
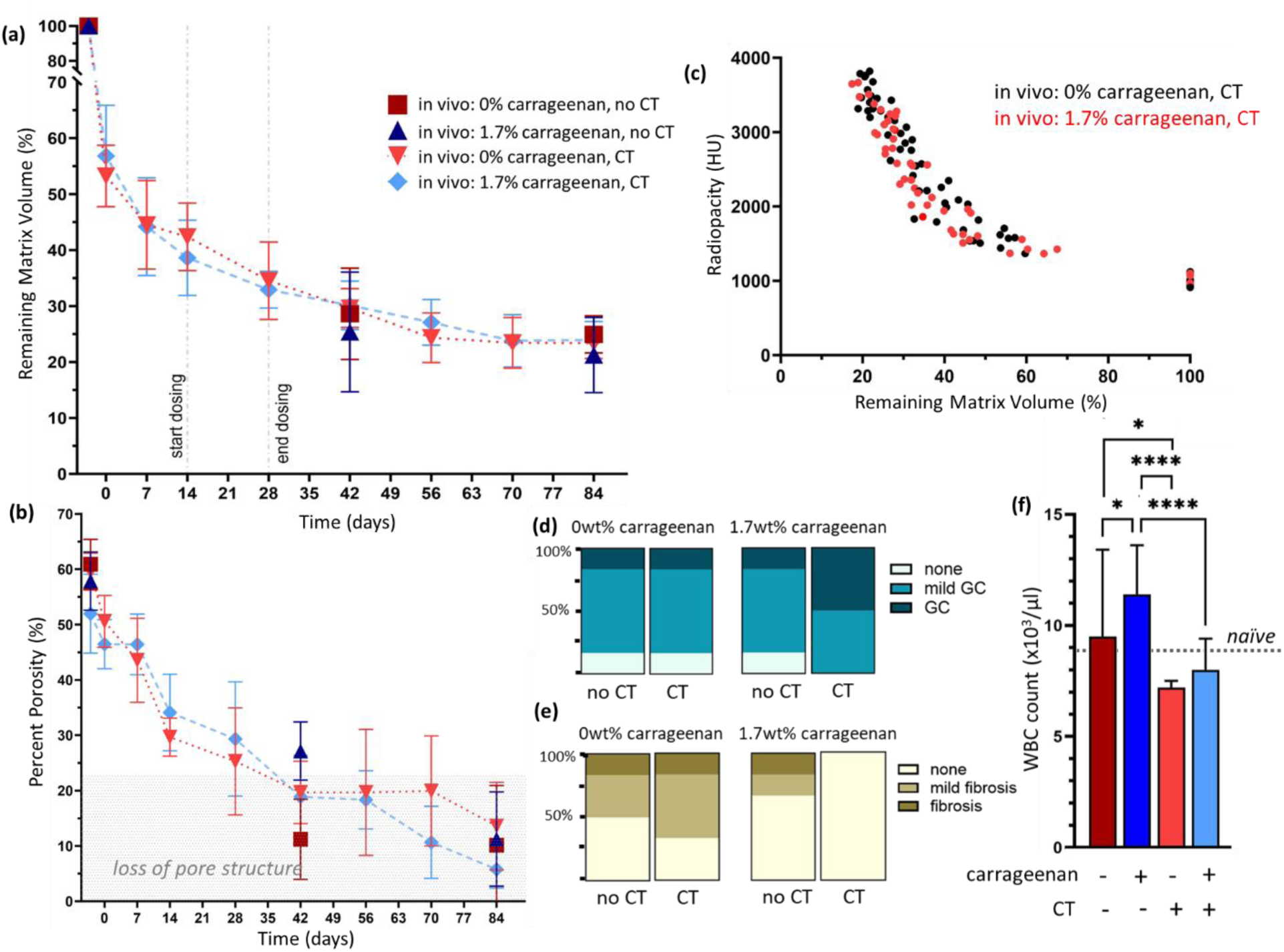
Serial monitoring of phantoms via CT did not significantly affect degradation kinetics and maintained inflammatory responses. (a) Serial imaging did not alter the rate of matrix volume loss but allowed for (b) longitudinal measurement of phantom features, such as percent porosity; below 25% porosity, a distinct pore structure was lost in all groups. During degradation, scaffolds collapsed and compacted, leading to (c) increased radiopacity as the volume of the matrix decreased. Radiation exposure did not significantly affect (d) the inflammatory response, reported as granulomatous cellulitis (GC). (e) Chronic inflammatory stimulation appeared to delay fibrosis in the tissues. (f) Systemically, x-ray exposure led to a lower white blood cell (WBC) count. *p < 0.05, **** p < 0.0001. All data shown as ± standard deviation.

Overall, local radiation exposure around the implants during imaging affected the circulating white blood cell count, but the total cell number was not significantly lower than in naive control animals, Figure 6(f). Histological examination of the tissue in direct contact with the phantoms showed no difference in the degree of inflammation response with CT monitoring either, Figure 6(d). Fibrosis was significantly lower with the induction of chronic inflammatory responses, Figure 6(e).

## 4 Discussion

Biomedical implantable devices often play critical roles in the stabilization of damaged tissues and support their regeneration. Whether designed to be permanent or to degrade over time, device impact on tissue repair is mediated by the immune system, particularly macrophages. Ideally, once acute inflammation responses subside, no fibrous encapsulation of the device occurs, leaving the porous structure free to be revascularized and infiltrated by native tissue. In this paradigm, the device serves as a support until native extra-cellular matrix can be produced to heal the damaged tissue. While all implantation will produce an acute inflammatory response due to surgery, over time that response should lessen; in disease states, inflammation persists, setting up an environment of chronic inflammation that does not allow tissue to heal.

Many properties affect the final inflammatory response to a biomedical device. For example, the material’s properties, like stiffness, can influence the foreign body reactions and local secretion of pro-inflammatory factors like TNFα due to macrophage activation via their cytoskeleton [19]. For polymeric phantoms, degradation products can also be a key influencer of long-term inflammation. Unfortunately, biomaterial degradation has been consistently difficult to predict from in vitro tests [7], and in the clinic extreme heterogeneity of patient populations further complicates the calculation of biomedical device lifetime. Therefore, a way to non-invasively track device degradation is required, which can transition into a clinical setting.

For this purpose, CT is an ideal imaging modality, as it is relatively low cost and high throughput [20]. A challenge for monitoring polymers with CT post-implantation is that they must be modified to introduce radiopacity for serial imaging or they are virtually invisible [10]. Tantalum oxide nanoparticles are ideal for imparting radiopacity as they are biocompatible, have a lower cost than noble metals like platinum or gold [21], and can be produced with a hydrophobic coating for easy homogeneous dispersion within synthetic polymers [8]. However, nanoparticle contrast agents must be validated as a method to track implants post-implantation, by confirming that the particles do not perturb inflammatory responses to the device and that serial monitoring does not significantly dampen normal immune responses due to radiation exposure [22].

### 4.1 Nanoparticle composites elicit a minor immune response

When polymer matrices are combined with contrast agents, not only polymer degradation products affect in vivo inflammatory responses, but the nanoparticle contrast agent released at the site of implantation can also affect macrophages. Therefore, the effects of nanoparticle addition were first studied in vitro with phantoms from PCL, PLGA 85:15 and PLGA 50:50, three biocompatible polymers covering the full range of predicted degradation behaviors from virtually non-degrading for PCL (years), slow degradation for PLGA 85:15 (months), and fast degradation PLGA 50:50 (weeks) [3,12]. The range of TaO_x_ incorporation was chosen to represent the baseline (0wt%), a minimum incorporation for radiopacity (5wt%) and the maximum shown biocompatible for primary cell culture (20wt%) [15].

The addition of nanoparticle contrast agents to PCL matrices had the least effect on macrophage secretion and protein expression. PCL is generally inert to macrophage degradation of the surface [23,24], but once implanted in vivo, is associated with IRF5 expression within CD68 macrophages at the implant site [24]. This was also observed in the current study, were PCL stimulated greater expression of IRF5 compared to PLGA films. In literature, IRF5 is associated with a pro-inflammatory phenotype, in particular with the initiation of angiogenesis in vivo [25,26].

PLGA substrates had consistently up-regulated secretion of the pro-inflammatory marker TNFα with nanoparticle incorporation. The addition of nanoparticles corresponds to increases in surface roughness [15]. While macrophage polarization is known to be sensitive to changes in topography and pore size in 3D structures, in general, macrophages are more sensitive to micron scale changes in topography, as opposed to nanoscale changes, such as those caused by the nanoparticles [27]. While smaller pores can lead to greater pro-inflammatory phenotypes [28], the nanoparticle addition did not alter the 3D porous structures in the present case. However, there is also evidence that PLGA degradation products, that affect cellular metabolism and local pH, can increase pro-inflammatory cytokine expression, particularly TNFα and IL-1B [1,29]. PLGA has a much faster degradation rate than PCL, with 85:15 degrading over months and 50:50 degrading in a matter of weeks, suggesting shorter time frames for stimulating inflammation. In addition, the degradation rate is further increased with nanoparticle addition [11]. The topographical changes to the PLGA surface seen via electron microscopy, suggest an even faster degradation profile for PLGA 50:50 than predicted from cell-free in vitro degradation studies.

In vivo, a healthy response to implanted materials requires initial pro-inflammatory activation (TNFα, IRF5) to stimulate angiogenesis at the region [24] and then transition to an anti-inflammatory phenotype (CCL17, TGFβ1) to induce matrix deposition [26]. Avoiding a fibrous encapsulation requires mediation by macrophages and is influenced by their phenotype, which can be a mixture of markers traditionally associated with pro- and anti-inflammatory activation [30]. For a biomaterial to stimulate regeneration as opposed to fibrous encapsulation, the device cannot bypass an inflammatory stage, but must encourage a timely transition of macrophage response towards the promotion of ECM production. The subtle changes in expression noted with the presence of TaO_x_ nanoparticles may, in fact, prove beneficial for the initial stimulation of angiogenesis.

### 4.2 Inflammation played a minor role on device degradation

The mouse model of chronic inflammation used in the current study was adapted from well known models that utilize carrageenan to induce an inflammatory response for testing anti-inflammatory drugs [31]. A chronic immune response was induced through repeated administration of carrageenan at the site of phantom implantation over two weeks. An inflammatory environment was created locally, based on histology, and systemically from the circulating white blood cell count. This was particularly evident at time points closer to the administration of carrageenan (week 6). Implantation of phantoms increased the inflammation response, both locally and systemically. The phantoms chosen for implantation consisted of a PLGA 50:50 matrix, the fastest degrading matrix tested in vitro, and one that is known to produce degradation products that further stimulate a pro-inflammatory environment [1,29]. In this way, the study represents the worst case scenario for the use of nanoparticle contrast agents to monitor device degradation post-implantation, one where the physiological environment is inherently unstable.

The inflammation response to implanted devices can be divided into an acute response, immediately after placement, and chronic longer-term responses, either triggered from infections or from the device degradation itself. Understanding inflammatory responses to implanted devices is often complicated by the surgery itself, which induces a separate acute immune response. While attempts have been made to describe degradation kinetics of polymers in vitro [2,3], there have been great discrepancies with in vivo work [32]. To better replicate degradation in vivo, studies have been conducted in a range of degradation buffers (pH 7.5-5.5), representing physiological environments ranging from healthy to lysosomal conditions [11]. Despite accelerated in vitro degradation conditions, the in vivo degradation of radiopaque phantoms occurred significantly faster in this study, particularly their structural collapse, Figure 5. This is likely due to the mechanical forces exerted by surrounding tissues, interstitial fluid flow, and the presence of proteases.

Another hypothesis for the faster degradation of devices in vivo has been an increased cellular response of macrophages, as inflammation pathways can be activated by changes in local pH such as those caused by the acidic degradation products of PLGA [33]. Despite the stimulation of a robust inflammatory response around implanted phantoms, there was no significant change to the measured degradation of the implants. However, the measure of degradation in this study was confined to quantification of polymer volume, indirectly measured via CT. Polymer degradation occurs in several stages, starting with the scission of the polymer chains, that affect their mechanical and chemical properties. Mass loss only occurs when polymer scission is severe enough to release monomers or short oligomers and thus lags behind loss of mechanical properties, which has been demonstrated in vitro [11].

### 4.3 Serial monitoring of devices does not affect physical processes

End-point analyses of device degradation has been the gold standard for tracking changes to device structure and tissue infiltration. However, the data that can be collected is necessarily limited to discrete time points spaced at arbitrary intervals and can suffer from a lack of power, given the number of animals required per group. On the other hand, serial in vivo monitoring post-implantation offers much more information on the intermediate stages of implant degradation. For example, the gold standard method for calculating volume remaining over-estimates the degradation rate, because it does not take into account the effect of tissue compression immediately post-implantation. Thus, by traditional methods, the results show that 80% of the matrix volume is lost by 12 weeks in vivo, when in reality, only 40% of the volume is lost. The collapse of the porous architecture of the devices occurred between weeks 4 and 6, and thus, mechanical stability would be lost at least by week 4 to allow macroscopic changes to the structure.

Serial monitoring is only useful if the act of imaging does not alter the test metrics significantly. In this study, we have successfully demonstrated that there is no significant change to degradation kinetics after exposure to radiation via CT imaging. There is also concern that the radiation dosage necessary to serially monitor fine features of implants can impair the animals’ inflammatory processes. In literature, the focus has been on minimizing the radiation dose per time point, but due to the complex nature of which tissues are exposed to radiation and what time course that exposure occurs in, biological effects are difficult to predict. With low radiation dose (16 mGy) mice have been scanned 3 times per week with no significant physiological changes to blood or litter size [34]. However, as radiation dose and scans per week increase significant decreases in circulating blood components, such as platelets and white blood cells, have been noted [22]. In the current study, changes in white blood cell count remained the most sensitive to CT exposure, but histology showed few significant changes to either granulomatous inflammation or fibrosis.

This study validates the use of TaO_x_ nanoparticle contrast agents for monitoring device degradation. There was low perturbation of macrophage response with the nanoparticle addition, just a mild increase in inflammation markers, likely mediated via increased rate of degradation from the device. Importantly, there was also no evidence of a burst release of nanoparticles from the polymer matrix post-implantation that could negatively impact the quality of the monitoring. While CT is valuable for tracking device information, dynamic biological responses like inflammation or neovascularization likely require a combination of imaging techniques such as optical imaging [35], positron emission tomography (PET) imaging [36], or MRI. The use of multiple imaging techniques is supported by the nanoparticles, as they did not interfere with other imaging techniques, such as MRI, Figure 6(f). The integration of preclinical and clinical monitoring techniques could revolutionize medical device development and ultimately provide easy markers for clinicians to personalize patient rehabilitation and avoid catastrophic failure of implantation.

## 5 Conclusion

A better understanding of how biomaterials interact with living tissue could usher in a new age of medical devices. However, this goal necessitates better methods for monitoring degradation and implant damage post-implantation. One method is through the functionalization of devices for medical imaging via contrast agent incorporation. Therefore, radiopaque TaO_x_ nanoparticles were added into biomedical polymer matrices homogeneously, without altering the structure of porous biomedical devices. With 5-20wt% TaO_x_ NPs, the devices could be radiographically distinguished from physiological tissue via CT monitoring and induced only a mild inflammatory reaction in the form of increased TNFα secretion, likely mediated through changes to the matrix properties. When radiopaque polymer devices were implanted in vivo, the stimulation of chronic inflammatory responses at the implant site did not significantly alter their degradation profile. Serial monitoring of radiopaque devices post-implantation was possible using CT, with no burst release of nanoparticles from the polymer matrix, and without dampening the native immune response below that of naive animals. Serial monitoring provided granular information on changes to the implanted devices, and could be combined with other medical imaging techniques to track dynamic biological changes, opening up new opportunities to engineer biological response through medical devices.

## 6 Acknowledgments

The authors would like to thank C. Mallet from the MSU Advanced Molecular Imaging Facility for obtaining the MRI images. This study was supported by the National Institute of Biomedical Imaging and Bioengineering of the NIH under award number R01EB029418. The content is solely the responsibility of the authors and does not necessarily represent the official views of the National Institutes of Health.

## References

[1] C. Maduka, M. Alhaj, E. Ural, M. Kuhnert, O. Habeeb, A. Schilmiller, K. Hankenson, S. Goodman, R. Narayan, and C. Contag. Stereochemistry determines immune cellular responses to polylactide implants. ACS Biomater. Sci. Eng. 2023; 9: 932–943.

[2] A. Goepferich. Mechanisms of polymer degradation and erosion. Biomaterials 1996; 17: 103–114.

[3] F. Alexis. Factors affecting the degradation and drug-release mechanism of poly(lactic acid) and poly(lactic acid)-co-(glycolic acid). Polymer International 2005; 54: 36–46.

[4] L. Huyer, S. Pascual-Gil, Y. Wang, S. Mandla, B. Yee, and M. Radisic. Advanced strategies for modulation of the material-macrophage interface. Adv. Funct. Mater. 2020; 30: 1909331.

[5] L. S. Saleh, L. D. Amer, B. J. Thompson, T. Danhorn, J. R. Knapp, S. L. Gibbings, S. Thomas, L. Barthel, B. P. O’Connor, W. J. Janssen, S. Alper, and S. J. Bryyant. Mapping macrophage polarization and origin during the progression of the foreign body response to a poly(ethylene glycol) hydrogel implant. Adv. Healthcare Mater. 2022; 11: 2102209.

[6] K. Ji, L. Mayernik, K. Moin, and B. F. Sloane. Acidosis and proteolysis in the tumor microenvironment. Cancer and Metastasis Reviews 2019; 38: 103–112.

[7] L. Lu, S. J. Peter, M. D. Lyman, H.-L. Lai, S. M. Leite, J. A. Tamada, S. Uyama, J. P. Vacanti, R. Langer, and A. G. Mikos. In vitro and in vivo degradation of porous poly(dl-lactic-co-glycolic acid) foams. Biomaterials 2000; 21: 1837–1845.

[8] S. Chakravarty, J. M. L. Hix, K. A. Wiewiora, M. C. Volk, E. Kenyon, D. D. Shuboni-Mulligan, B. Blanco-Fernandez, M. Kiupel, J. Thomas, L. F. Sempere, and E. M. Shapiro. Tantalum oxide nanoparticles as versatile contrast agents for x-ray computed tomography. Nanoscale 2020; 12: 7720–7734.

[9] J. M. Crowder, N. Bates, J. Roberts, A. S. Torres, and P. J. Bonitatibus. Determination of tantalum from tantalum oxide nanoparticle x-ray/ct contrast agents in rat tissues and bodily fluids by ICP-OES. J. Anal. At. Spectrom 2016; 31: 1311–1317.

[10] K. Pawelec, S. Chakravarthy, J. Hix, K. Perry, L. van Holsbeeck, R. Fajardo, and E. Shapiro. Design consideration to facilitate clinical radiological evaluation of implantable biomedical structures. ACS Biomaterials Sci. Eng. 2021; 7(2): 718–726.

[11] K. Pawelec, E. Tu, S. Chakravarty, J. Hix, L. Buchanan, L. Kenney, F. Buchanan, N. Chatterjee, S. Das, A. Alessio, and E. Shapiro. Incorporating tantalum oxide nanoparticles into implantable polymeric biomedical devices for radiological monitoring. Adv. Healthcare Mater. 2023: 2203167.

[12] M. Bartnikowski, T. R. Dargaville, S. Ivanovski, and D. W. Hutmacher. Degradation mechanisms of polycaprolactone in the context of chemistry, geometry and environment. Progress in Polym. Sci. 2019; 96: 1–20.

[13] G. Narayanan, V. N. Vernekar, E. L. Kuyinu, and C. T. Laurencin. Poly(lactic acid)-based biomaterials for orthopaedic regenerative engineering. Adv. Drug Delivery Reviews 2016; 107: 247–276.

[14] I. Pineda-Torra, M. Gage, A. de Juan, and O. M. Pello. Isolation, culture and polarization of murine bone marrow-derived and peritoneal macrophages, vol. 1339, ch. 6, pp. 101–110. Springer.

[15] K. Pawelec, J. Hix, and E. Shapiro. Functional attachment of primary neurons and glia on radiopaque implantable biomaterials for nerve repair. Nanomedicine: Nanotechnology, Biology, and Medicine 2023; 52: 102692.

[16] J. Meganck and B. Liu. Dosimetry in micro-computed tomography: a review of the measurement methods, impacts, and characterization of the quantum GX. Mol Imaging Biol 2017; 19: 499–511.

[17] P. A. Yushkevich, J. Piven, H. C. Hazlett, R. G. Smith, S. Ho, J. C. Gee, and G. Gerig. User-guided 3D active contour segmentation of anatomical structures: Significantly improved efficiency and reliability. NeuroImage 2006; 31: 1116–1128.

[18] K. Spiller, E. Wrona, S. Romero-Torres, I. Pallotta, P. Graney, C. Witherel, L. Panicker, R. Feldman, A. Urbanska, L. Santambrogio, G. Vunjak-Novakovic, and D. Freytes. Differential gene expression in human, murine, and cell line-derived macrophages upon polarization. Experimental Cell Research 2016; 347: 1–13.

[19] Y. Ni, H. Qi, F. Zhang, S. Jiang, Q. Tang, W. Cai, W. Mo, R. Miron, and Y. Zhang. Macrophages modulate stiffness-related foreign body responses through plasma membrane deformation. PNAS 2022; 120(3): e2213837120.

[20] J. Wallyn, N. Anton, S. Akram, and T. F. Vandamme. Biomedical imaging: principles, technologies, clinical aspects, contrast agents, limitations and future trends in nanomedicines. Pharm Res 2019;36: 78.

[21] K. R. Sneha and G. S. Sailaja. Intrinsically radiopaque biomaterial assortments: a short review on the physical principles, x-ray imageability, and state-of-the-art developments. J. Mater. Chem. B 2021; 9: 8569.

[22] N. Berghen, K. Dekoster, E. Marien, D. J, H. A, J. Wouters, J. Deferme, T. Vosselman, E. Tiest, M. Lox, J. Vanoirbeek, E. De Langhe, R. Bogaerts, M. Hoylaerts, R. Lories, and G. Vande Velde. Radiosafe micro-computed tomography for longitudinal evaluation of murine disease models. Sci. Rep. 2019; 9: 17598.

[23] E. Bat, T. van Kooten, J. Feijen, and D. W. Grijpma. Macrophage-mediated erosion of gamma irradiated poly(trimethylene carbonate) films. Biomaterials 2009; 30: 3652–3661.

[24] E. Dondossola, B. M. Holzapfel, S. Alexander, S. Filippini, D. W. Hutmacher, and P. Friedl. Examination of the foreign body response to biomaterials by nonlinear intravital microscopy. Nature Biomedical Engineering 2016; 1: 0007.

[25] H. Almuttaqi and I. A. Udalova. Advances and challenges in targeting IRF5, a key regulator of inflammation. FEBS Journal 2018; 286: 1624–1637.

[26] C. E. Witherel, D. Abebayehu, T. H. Barker, and K. L. Spiller. Macrophage and fibroblast interactions in biomaterial-mediated fibrosis. Adv Healthc Mater 2019; 8(4): e1801451.

[27] S. Chen, J. A. Jones, Y. Xu, H.-Y. Low, J. M. Anderson, and K. W. Leong. Characterization of topographical effects on macrophage behavior in a foreign body response model. Biomaterials 2010; 31: 3479–3491.

[28] E. M. Sussman, M. C. Halpin, J. Muster, R. T. Moon, and B. D. Ratner. Porous implants modulate healing and induced shifts in local macrophage polarization in the foreign body reaction. Annals of Biomedical Eng. 2014; 42(7): 1508–1516.

[29] S. Ma, X. Feng, F. Liu, B. Wang, H. Zhang, and X. Niu. The pro-inflammatory response of macrophages regulated by acid degradation products of poly(lactide-co-glycolide) nanoparticles. Engineering in Life Sciences 2021; 21: 709–720.

[30] C. E. Witherel, K. Sao, B. K. Brisson, B. Han, S. W. Volk, R. J. Petrie, L. Han, and K. L. Spiller. Regulation of extracellular matrix assembly and structure by hybrid M1/N2 macrophages. Biomaterials 2021; 269: 120667.

[31] J. C. Fehrenbacher and K. E. McCarson. Models of inflammation: carrageenan air pouch. Current Protocols 2021; 1: e183.

[32] K. Fu, D. W. Pack, A. M. Kibanov, and R. Langer. Visual evidence of acidic environment within degrading poly(lactic-co-glycolic acid) (PLGA) microspheres. Pharmaceutical Res. 2000; 17(1): 100–106.

[33] A. Riemann, H. Wussling, H. Loppnow, H. Fu, S. Reime, and O. Thews. Acidosis differently modulates the inflammatory program in monocytes and macrophages. Biochimica et Biophysica Acta 2016; 1862: 72–81.

[34] N. Miyahara, T. Kokubo, Y. Hara, A. Yamada, T. Koike, and Y. Arai. Evaluation of X-ray doses and their corresponding biological effects on experimental animals in cone-beam micro-CT scans (r-mct2). Radiol Phys Technol 2016; 9: 60–68.

[35] H. Scales, M. Ierna, K. Smith, K. ross, G. Meiklejohn, J. Patterson-Kane, I. McInnes, J. Brewer, P. Garside, and P. Maffia. Assessment of murine collagen-induced arthritis by longitudinal non-invasive duplexed molecular optical imaging. Rheumatology 2016; 55: 564–572.

[36] C. Tondera, S. Hauser, A. Krueger-Genge, F. Jung, A. T. Neffe, L. A., R. Klopfleisch, J. Steinbach, C. Neuber, and J. Pietzsch. Gelatin-based hydrogel degradation and tissue interaction in vivo: Insights from multimodal preclinical imaging in immunocompetent nude mice. Theranostics 2016; 6: 2114–2128.

