## Supplemental for "In vivo Biomedical Imaging of Immune Tolerant, Radiopaque Nanoparticle-Embedded Polymeric Device Degradation"

#### S1. Macrophage Polarization

As a control for the ability of the isolated mouse bone marrow derived macrophages (BMDM) to polarize, 2-3×$10^4$ cells/well were plated on tissue culture plastic and stimulated with soluble factors. Two days prior to harvesting supernatant or protein, control cells were treated with media supplemented with polarization factors, to serve as positive controls for assessing polarization on film surfaces. The three control groups were 1) no polarization (M0), using unsupplemented differentiation media, 2) pro-inflammatory M(LPS/IFNγ) (differentiation media supplemented with 100 ng/ml lipopolysaccharide (LPS, Sigma, L4391) and 50 ng/ml interferon gamma (IFNγ, Pepro Tech, 315-05-20UG)) or 3) anti-inflammatory M(IL4/IL13) (differentiation media supplemented with 40 ng/ml IL-4 (Pepro Tech, 214-14-5UG) and 20 ng/ml IL13 (Pepro Tech 210-13-2UG). All tests were done in triplicate from different cell lots, with two technical replicates each, and presented as mean ± standard error.

At the time of harvest (24-48 hrs post polarization), supernatant was probed for tumor necrosis factor alpha (TNFα, R&D Systems, mta006), IL10 (Invitrogen, BMS614INST), and CCL17 (R&D Systems, mcc170). On day 7, cellular levels of arginase I were also quantified by ELISA (abcam, ab269541), lysing cells in protein lysis buffer for 15 minutes on ice with the reagents provided by the kit, following manufacturer's instructions. Arginase-1 concentration was normalized to protein loading. In addition, the nitrite concentration in the supernatant was measured via the Griess assay (Promega, TB229), per manufacturer's instructions. In addition, protein expression of IRF5, TGFβ1, and integrin $\alpha_V$ were quantified via western blotting (Section S.3). The changes in polarization markers are presented in **Figure S1**.

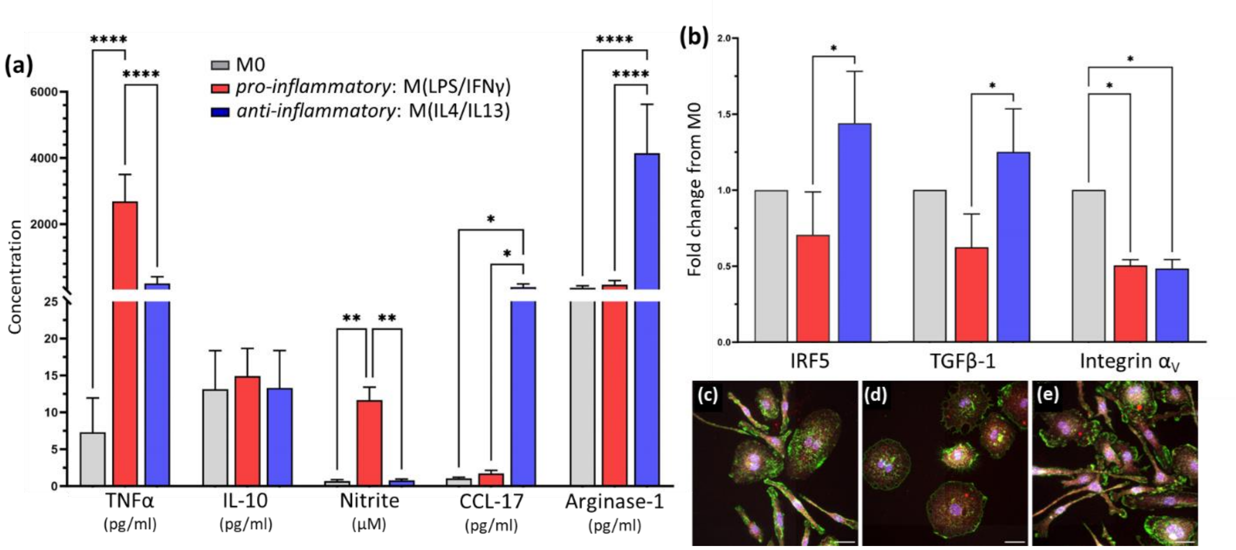


**Figure S1**. Mouse BMDMs could be polarized to a pro-inflammatory M(LPS/IFNγ) phenotype or anit-inflammatory M(IL4/IL13) phenotype. (a) Compared to naïve BMDM, supernatant concentration of TNFα and nitrite was significantly higher for M(LPS/IFNγ), while M(IL4/IL13) cells secreted significantly more CCL-17 and Arginase-1 expression; the unit of concentration for each factor is noted below the factor in parenthesis. (b) Protein expression was significantly affected during polarization; all protein expression was normalized to naïve BMDM expression. Changes in cell morphology were also noted with BMDM polarization: (c) M0 cells, (d) M(LPS/IFNγ) cells and (e) M(IL4/IL13) cells; actin cytoskeleton (green), IRF5 (white), CD68 (red), and nucleus (DAPI, blue). Scale bar (c-e): 25µm.

### S2. Macrophages on Polymer Substrates

Scanning electron microscopy micrographs were taken of films after 1 week of culture and compared to films that were incubated at 37°C in culture media without cells. Films were composed of either poly(caprolactone) (PCL), or poly(lactide co-glycolide) (PLGA) 85:15 or 50:50. To retain cell morphology, fixed samples were dried in serial ethanol dilutions, at least 1 hour incubation in each: 70% ethanol, 80% ethanol, 90% ethanol, 100% ethanol. Finally, samples were incubated in hexamethyldi-silazane (Sigma) for 10 minutes and allowed to air dry. Dried samples were adhered to 13mm aluminum stubs and sputter coated with platinum. Surfaces were examined using a Zeiss Auriga, at 2 keV in scanning electron mode. High-magnification images of film surfaces with and without BMDMs is presented in **Figure S2**.

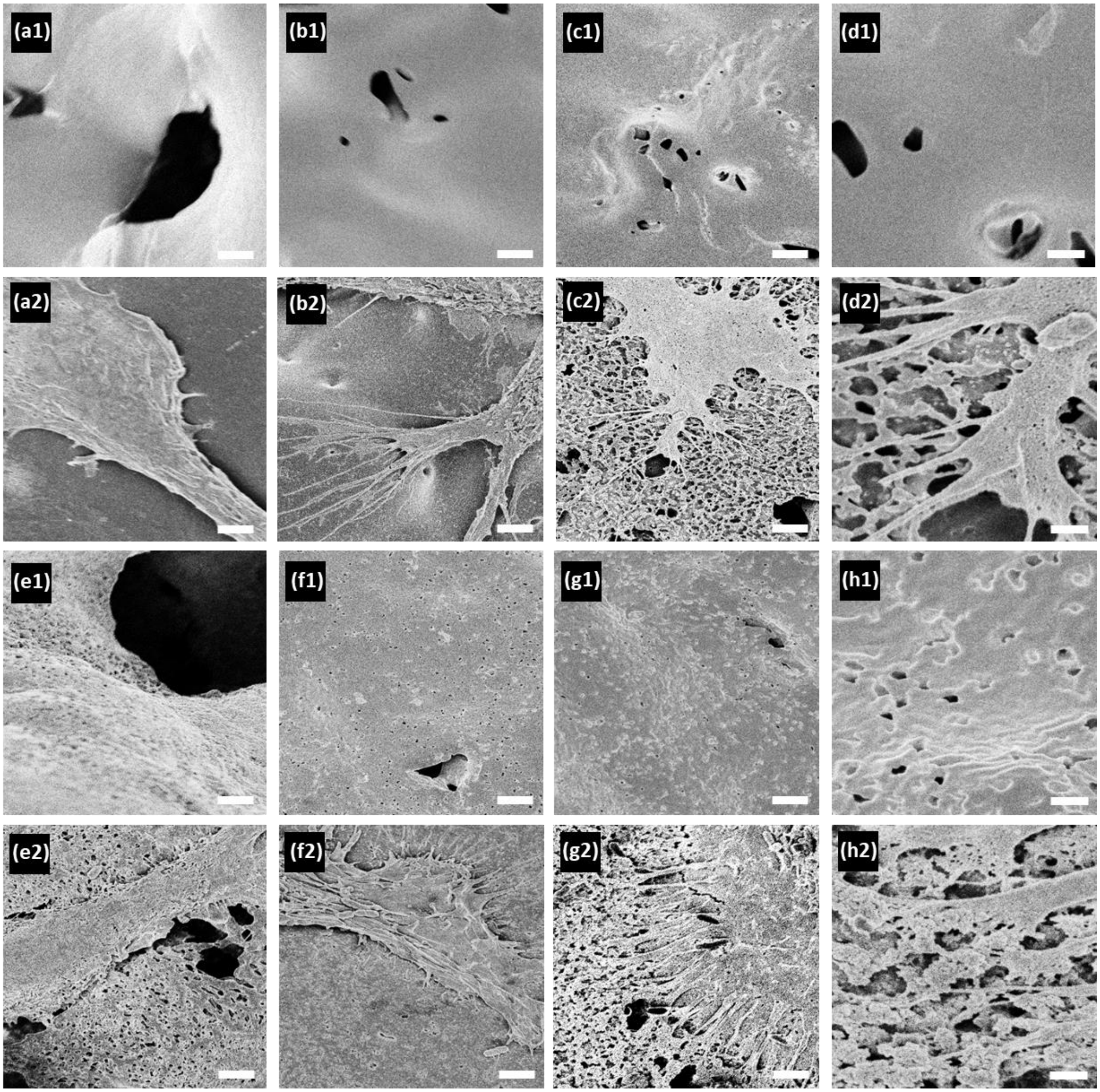


**Figure S2**. Polymer films with and without nanoparticles could be degraded by mouse BMDMs over one week in culture, dictated by the polymer matrix and not the addition of nanoparticles. Films without nanoparticle (0wt% $TaO_x$) were imaged (1) without cells and (2) after cell attachment: (a) PCL, (b) PLGA 85:15, (c-d) PLGA 50:50. With 20wt% $TaO_x$ nanoparticle incorporation, the same trends were noted (1) without cells and (2) with cell attachment: (e) PCL, (f) PLGA 85:15, (g-h) PLGA 50:50. Only PLGA 50:50 showed degradation, mediated by the BMDMs. Scale (a-c, e-g): 2 µm, (d, h): 500 nm.

**S.3 Protein Expression**

Western blotting was used to quantify protein expression. BMDMs cultured on films were lysed in Ripa buffer (supplemented with phosphatase inhibitors and protease inhibitors), and spun down at 15,000×g for 5 min at 4 °C. After which, supernatants were collected and the protein concentration of the supernatant was measured (BioRad, BCA kit). Supernatants were combined with 2× Laemmli buffer, incubated at 60°F for 20 min, separated by SDS/ PAGE (2-4 μg of protein per lane), and transferred to PVDF membranes. Transfer was accomplished using the iBlot 2 Gel Transfer Device (invitrogen, IB21001), a dry transfer system, at the recommended setting of 20 V for 1 min, 23 V for 4 min and 25 V for 2 minutes (7 min total). PVDF membranes were blocked with 5% dried milk in PBST (phosphate buffered saline pH 7.4, containing 0.1% Tween-20) and probed with primary antibodies specific for IRF5 (1:875, abcam, ab181553), TGFβ1 (1:700, abcam, ab215715), or integrin αV(1:1100-1:1400, abcam, ab302640) at 4 °C overnight. The next day, the membrane was washed and incubated with anti-mouse IgG-HRP (12-349, Millipore) or anti-rabbit IgG-HRP (ab205718, abcam) at room temperature for 2 hr and further washed in PBST six times.

Protein bands were visualized with ECL Plus substrate (cytiva, RPN2232) on a Li-Cor Odyssey FC and resulting signal was quantified via Image J. Membranes were then washed in PBS and stripped for 15 minutes in Restore Western Blot Stripping Buffer (Thermo Scientific 21059) with subsequent PBS washing. Membranes were either blocked and reprobed or were stained for total protein using the BLOT-Fast Stain (G Biosciences, cat# 786-34) according to manufacturer's directions. The total protein signal was imaged under visible light on a c300 imager (azure biosystems).

Signal for total protein in each lane was quantified on Image J, and the protein loading was normalized to the signal from the first lane. The normalized signal was used to adjust the measured band intensity from antibody staining, also quantified using Image J. For comparison of nanoparticle effect, all protein expression was recorded as a fold change from matrices with 0wt% $TaO_x$. BMDM polarization for control cells not cultured on films was normalized to the expression of non-stimulated macrophages (M0). The final protein quantification reported is the result of three biological replicates (2 technical replicates each). Samples of PCL + 0wt% $TaO_x$ were present on all blots and used to normalize differences between membrane exposure.

In the following figures (**Figures S3-S8**), each set of membranes is presented with all protein bands shown, along with the total protein staining for each membrane. Where no marker was present on the blot, the molecular weight of protein bands was determined from an initial blot, with a molecular weight marker present.

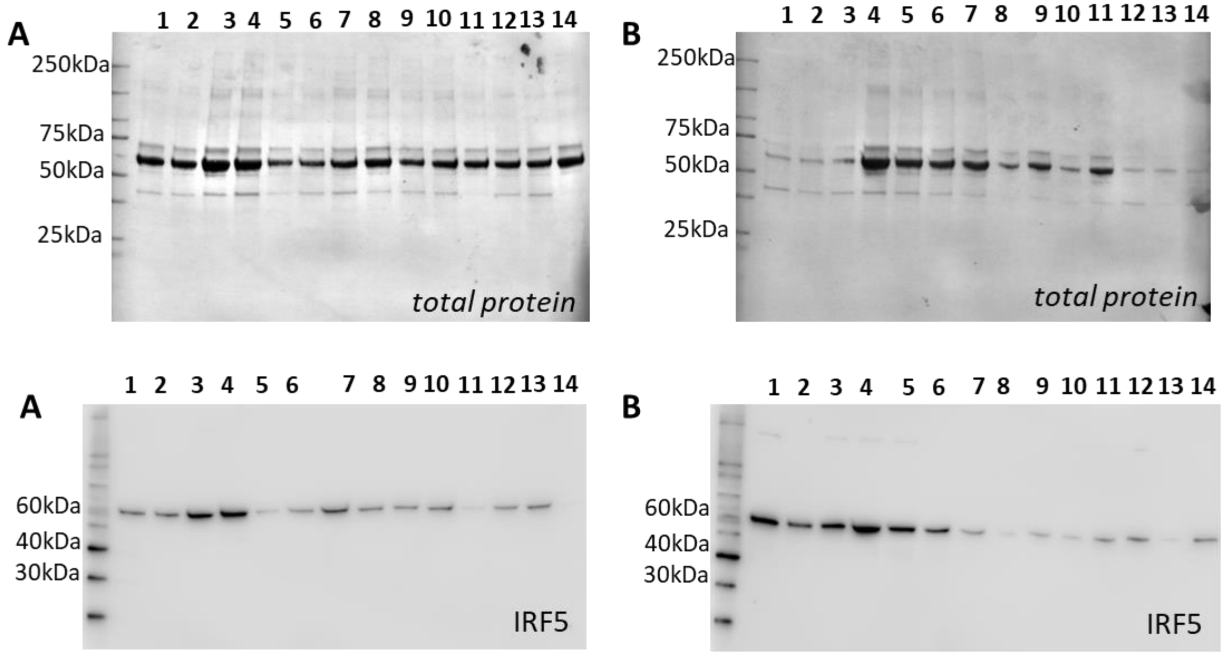


**A**

**1:** PLGA 50:50 (0wt% $TaO_x$)
**2:** PLGA 85:15 (0wt% $TaO_x$)
**3:** PCL (0wt% $TaO_x$)
**4:** PCL (5wt% $TaO_x$)
**5:** PCL (20wt% $TaO_x$)
**6:** PCL (20wt% $TaO_x$)
**7:** PCL (5wt% $TaO_x$)
**8:** PCL (0wt% $TaO_x$)
**9:** PLGA 50:50 (0wt% $TaO_x$)
**10:** PLGA 85:15 (0wt% $TaO_x$)
**11:** PLGA 85:15 (5wt% $TaO_x$)
**12:** PLGA 85:15 (20wt% $TaO_x$)
**13:** PLGA 85:15 (20wt% $TaO_x$)
**14:** PLGA 85:15 (5wt% $TaO_x$)

**B**

**1:** M0
**2:** M(LPS/IFNγ)
**3:** M(IL4/IL13)
**4:** PCL (0wt% $TaO_x$)
**5:** PLGA 50:50 (0wt% $TaO_x$)
**6:** PLGA 50:50 (5wt% $TaO_x$)
**7:** PLGA 50:50 (20wt% $TaO_x$)
**8:** PLGA 50:50 (20wt% $TaO_x$)
**9:** PLGA 50:50 (5wt% $TaO_x$)
**10:** PLGA 50:50 (0wt% $TaO_x$)
**11:** PCL(5wt% $TaO_x$)
**12:** M0
**13:** M(LPS/IFNγ)
**14:** M(IL4/IL13)

**Figure S3**: Western blots of IRF5 expression (biological replicate 1). Top row: total protein, second row: IRF5 staining, third row: sample labels.

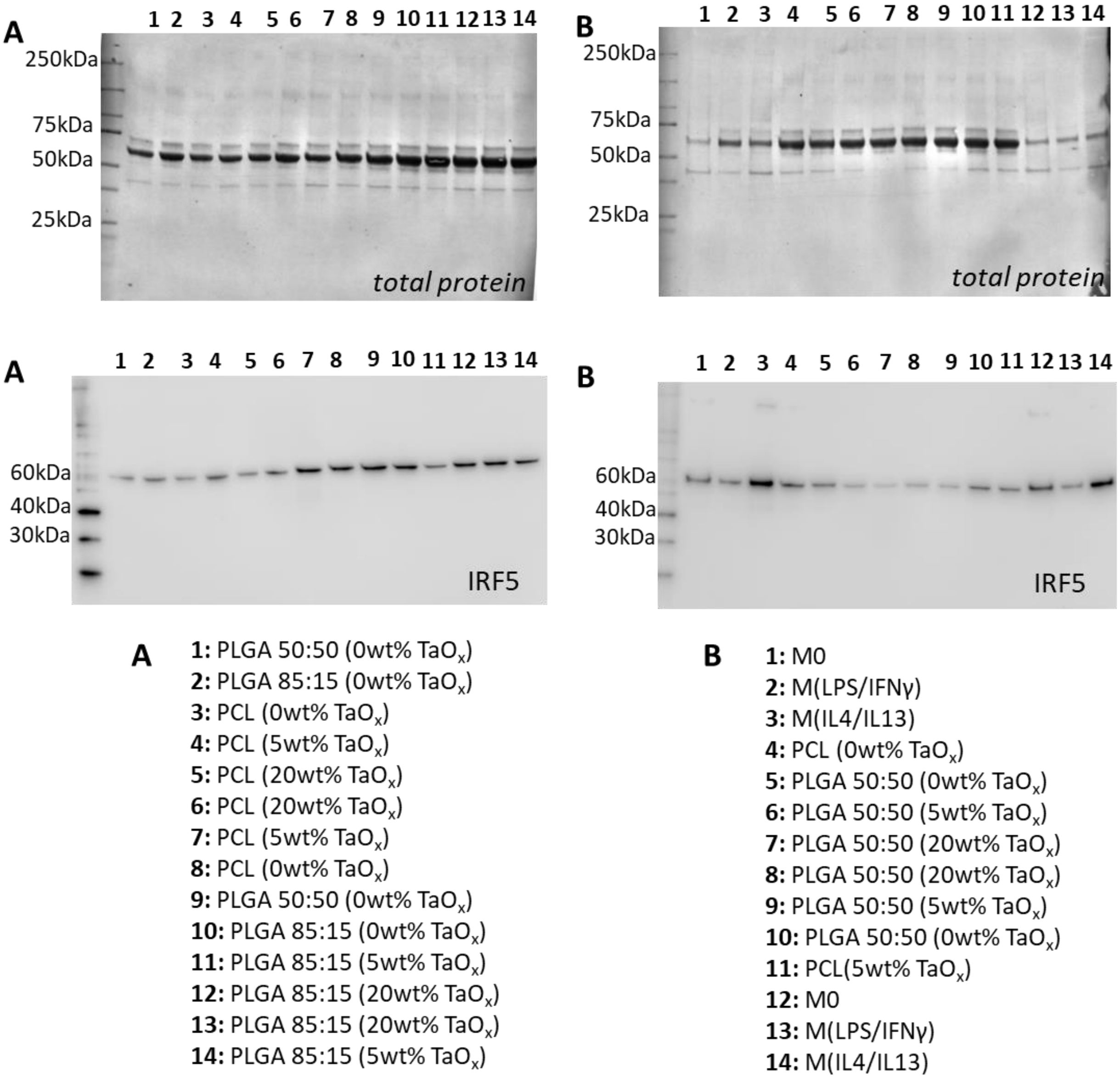


**Figure S4**: Western blots of IRF5 expression (biological replicate 2). Top row: total protein, second row: IRF5 staining, third row: sample labels.

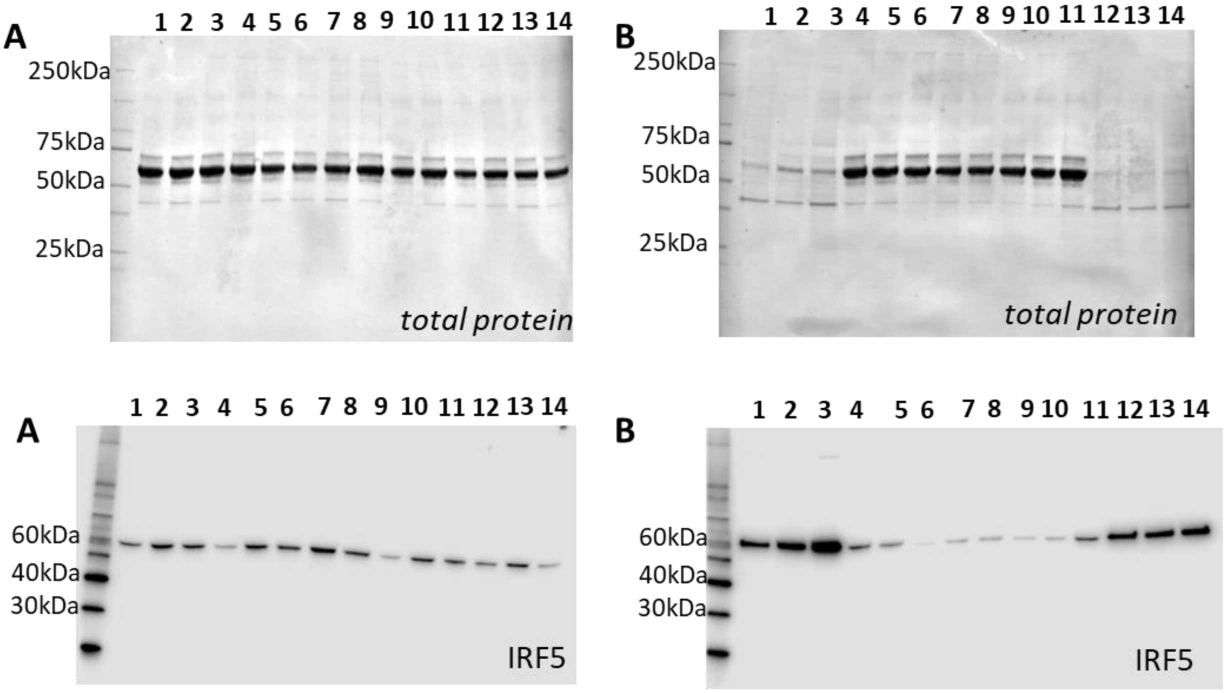


**A**

**1:** PLGA 50:50 (0wt% $TaO_x$)
**2:** PLGA 85:15 (0wt% $TaO_x$)
**3:** PCL (0wt% $TaO_x$)
**4:** PCL (5wt% $TaO_x$)
**5:** PCL (20wt% $TaO_x$)
**6:** PCL (20wt% $TaO_x$)
**7:** PCL (5wt% $TaO_x$)
**8:** PCL (0wt% $TaO_x$)
**9:** PLGA 50:50 (0wt% $TaO_x$)
**10:** PLGA 85:15 (0wt% $TaO_x$)
**11:** PLGA 85:15 (5wt% $TaO_x$)
**12:** PLGA 85:15 (20wt% $TaO_x$)
**13:** PLGA 85:15 (20wt% $TaO_x$)
**14:** PLGA 85:15 (5wt% $TaO_x$)

**B**

**1:** M0
**2:** M(LPS/IFNγ)
**3:** M(IL4/IL13)
**4:** PCL (0wt% $TaO_x$)
**5:** PLGA 50:50 (0wt% $TaO_x$)
**6:** PLGA 50:50 (5wt% $TaO_x$)
**7:** PLGA 50:50 (20wt% $TaO_x$)
**8:** PLGA 50:50 (20wt% $TaO_x$)
**9:** PLGA 50:50 (5wt% $TaO_x$)
**10:** PLGA 50:50 (0wt% $TaO_x$)
**11:** PCL(5wt% $TaO_x$)
**12:** M0
**13:** M(LPS/IFNγ)
**14:** M(IL4/IL13)

**Figure S5**: Western blots of IRF5 expression (biological replicate 3). Top row: total protein, second row: IRF5 staining, third row: sample labels.

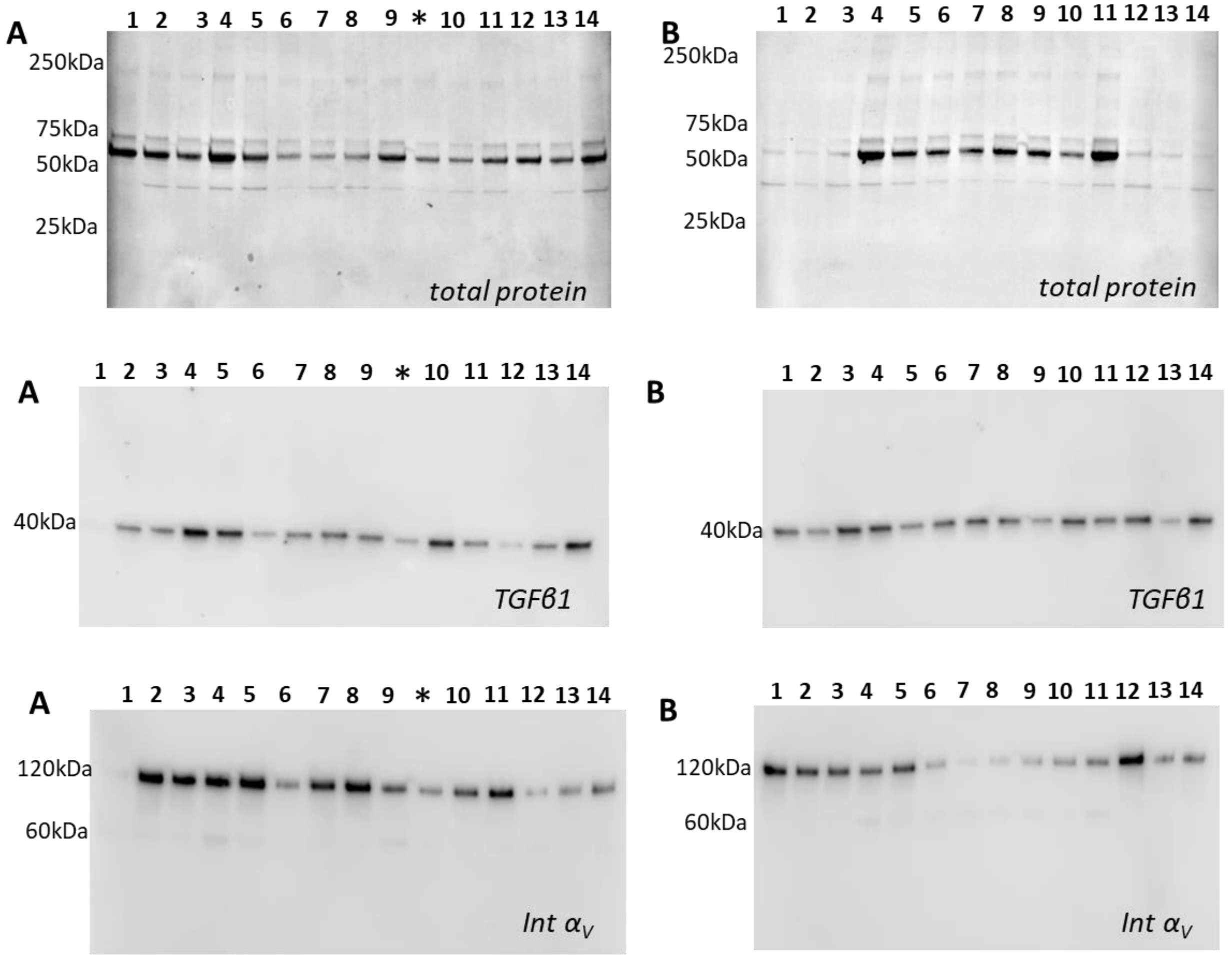


*incorrect loading

**A**

**1:** PLGA 85:15 (5wt% $TaO_x$)
**2:** PLGA 50:50 (0wt% $TaO_x$)
**3:** PLGA 85:15 (0wt% $TaO_x$)
**4:** PCL (0wt% $TaO_x$)
**5:** PCL (5wt% $TaO_x$)
**6:** PCL (20wt% $TaO_x$)
**7:** PCL (20wt% $TaO_x$)
**8:** PCL (5wt% $TaO_x$)
**9:** PCL (0wt% $TaO_x$)
**10:** PLGA 50:50 (0wt% $TaO_x$)
**11:** PLGA 85:15 (0wt% $TaO_x$)
**12:** PLGA 85:15 (5wt% $TaO_x$)
**13:** PLGA 85:15 (20wt% $TaO_x$)
**14:** PLGA 85:15 (20wt% $TaO_x$)

**B**

**1:** M0
**2:** M(LPS/IFNγ)
**3:** M(IL4/IL13)
**4:** PCL (0wt% $TaO_x$)
**5:** PLGA 50:50 (0wt% $TaO_x$)
**6:** PLGA 50:50 (5wt% $TaO_x$)
**7:** PLGA 50:50 (20wt% $TaO_x$)
**8:** PLGA 50:50 (20wt% $TaO_x$)
**9:** PLGA 50:50 (5wt% $TaO_x$)
**10:** PLGA 50:50 (0wt% $TaO_x$)
**11:** PCL(5wt% $TaO_x$)
**12:** M0
**13:** M(LPS/IFNγ)
**14:** M(IL4/IL13)

**Figure S6**: Western blots of TGFβ1 and integrin $\alpha_V$ expression (biological replicate 1). Top row: total protein, second row: TGFβ1 staining, third row: integrin $\alpha_V$ staining, forth row: sample labels. *indicates a lane that was loaded with an incorrect amount of protein.

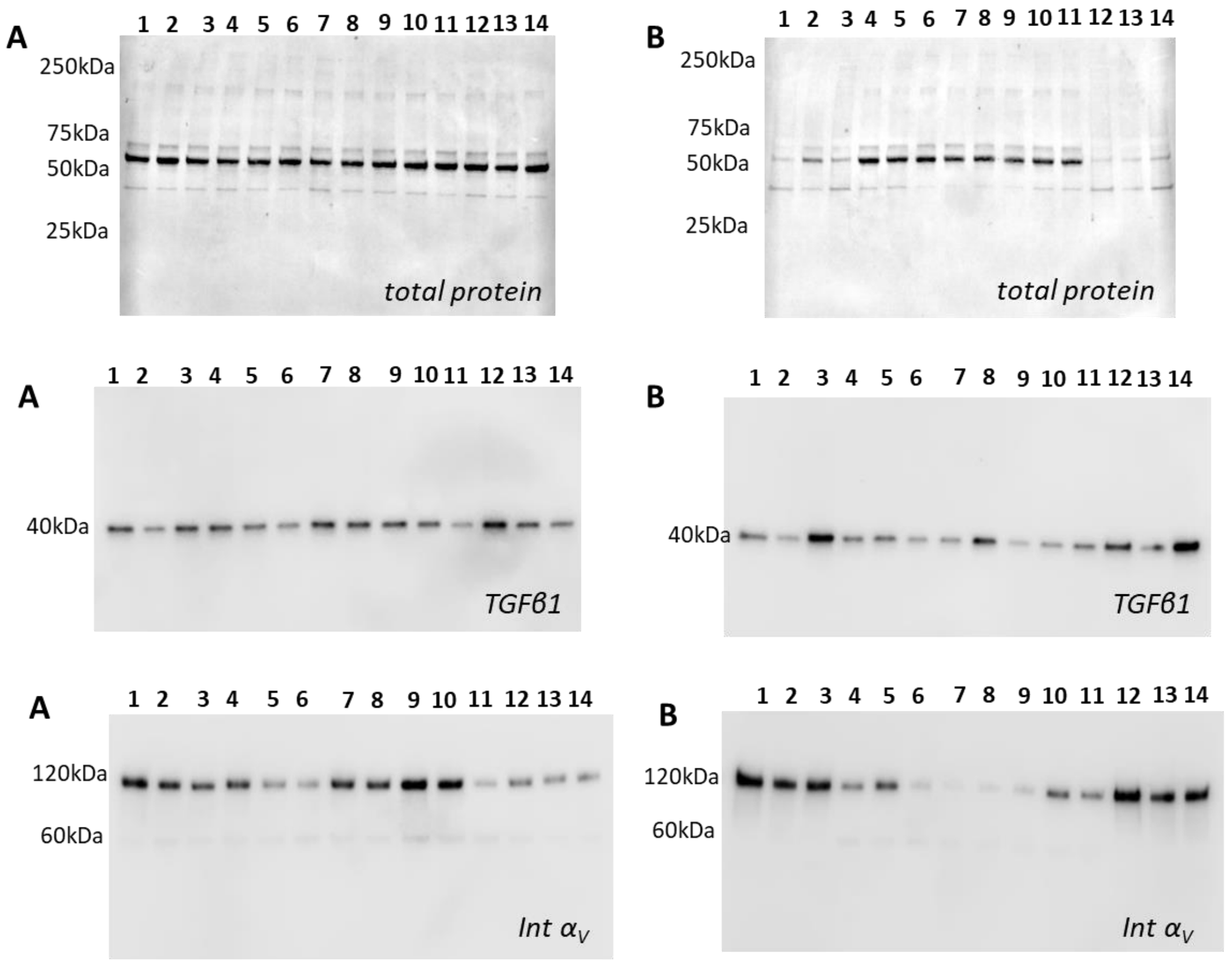


**A**

**1:** PLGA 50:50 (0wt% $TaO_x$)
**2:** PLGA 85:15 (0wt% $TaO_x$)
**3:** PCL (0wt% $TaO_x$)
**4:** PCL (5wt% $TaO_x$)
**5:** PCL (20wt% $TaO_x$)
**6:** PCL (20wt% $TaO_x$)
**7:** PCL (5wt% $TaO_x$)
**8:** PCL (0wt% $TaO_x$)
**9:** PLGA 50:50 (0wt% $TaO_x$)
**10:** PLGA 85:15 (0wt% $TaO_x$)
**11:** PLGA 85:15 (5wt% $TaO_x$)
**12:** PLGA 85:15 (20wt% $TaO_x$)
**13:** PLGA 85:15 (20wt% $TaO_x$)
**14:** PLGA 85:15 (5wt% $TaO_x$)

**B**

**1:** M0
**2:** M(LPS/IFNγ)
**3:** M(IL4/IL13)
**4:** PCL (0wt% $TaO_x$)
**5:** PLGA 50:50 (0wt% $TaO_x$)
**6:** PLGA 50:50 (5wt% $TaO_x$)
**7:** PLGA 50:50 (20wt% $TaO_x$)
**8:** PLGA 50:50 (20wt% $TaO_x$)
**9:** PLGA 50:50 (5wt% $TaO_x$)
**10:** PLGA 50:50 (0wt% $TaO_x$)
**11:** PCL(5wt% $TaO_x$)
**12:** M0
**13:** M(LPS/IFNγ)
**14:** M(IL4/IL13)

**Figure S7**: Western blots of TGFβ1 and integrin $\alpha_V$ expression (biological replicate 2). Top row: total protein, second row: TGFβ1 staining, third row: integrin $\alpha_V$ staining, forth row: sample labels.

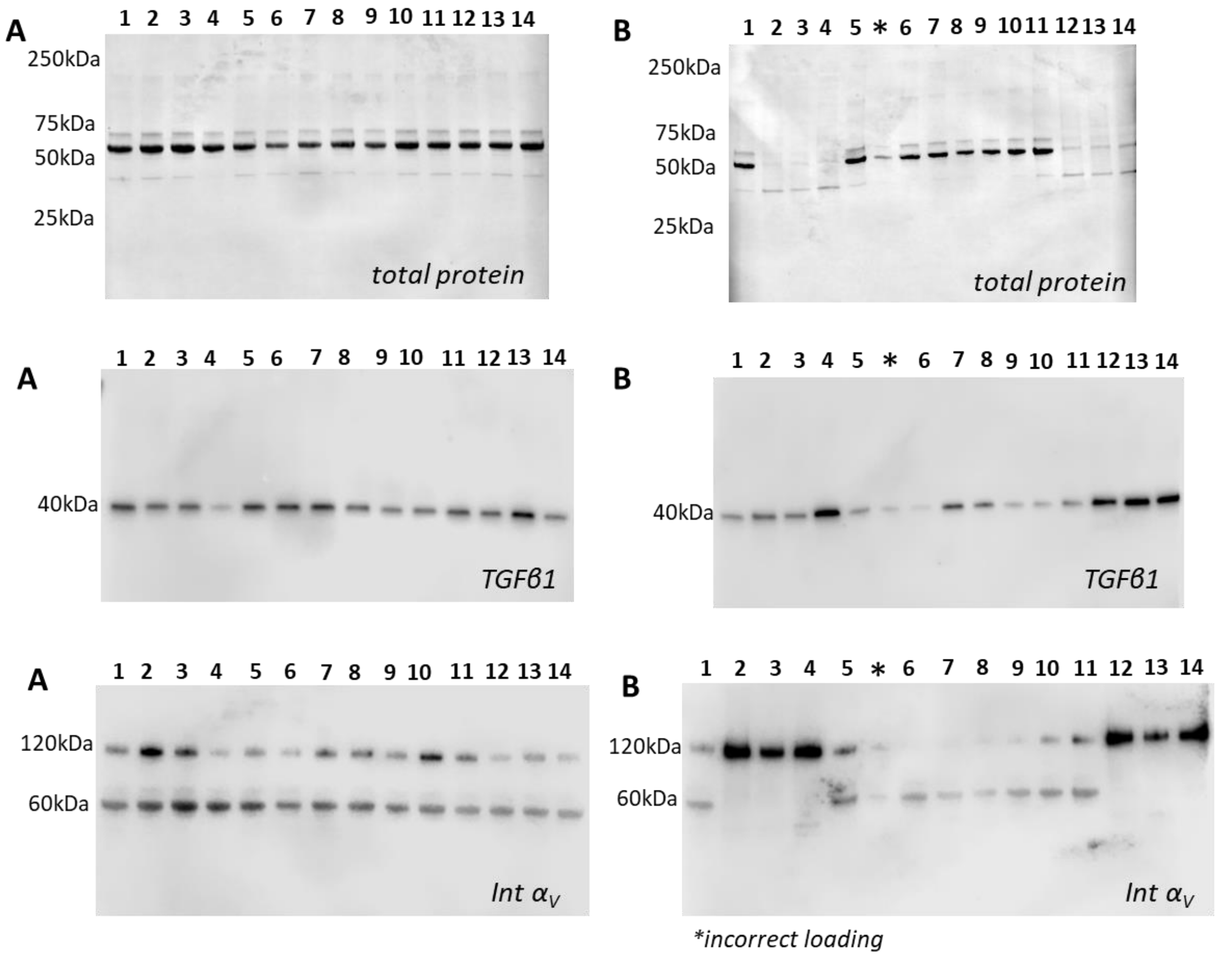


**A**

**1:** PLGA 50:50 (0wt% $TaO_x$)
**2:** PLGA 85:15 (0wt% $TaO_x$)
**3:** PCL (0wt% $TaO_x$)
**4:** PCL (5wt% $TaO_x$)
**5:** PCL (20wt% $TaO_x$)
**6:** PCL (20wt% $TaO_x$)
**7:** PCL (5wt% $TaO_x$)
**8:** PCL (0wt% $TaO_x$)
**9:** PLGA 50:50 (0wt% $TaO_x$)
**10:** PLGA 85:15 (0wt% $TaO_x$)
**11:** PLGA 85:15 (5wt% $TaO_x$)
**12:** PLGA 85:15 (20wt% $TaO_x$)
**13:** PLGA 85:15 (20wt% $TaO_x$)
**14:** PLGA 85:15 (5wt% $TaO_x$)

**B**

**1:** PLGA 50:50 (0wt% $TaO_x$)
**2:** M0
**3:** M(LPS/IFNγ)
**4:** M(IL4/IL13)
**5:** PCL (0wt% $TaO_x$)
**6:** PLGA 50:50 (5wt% $TaO_x$)
**7:** PLGA 50:50 (20wt% $TaO_x$)
**8:** PLGA 50:50 (20wt% $TaO_x$)
**9:** PLGA 50:50 (5wt% $TaO_x$)
**10:** PLGA 50:50 (0wt% $TaO_x$)
**11:** PCL(5wt% $TaO_x$)
**12:** M0
**13:** M(LPS/IFNγ)
**14:** M(IL4/IL13)

**Figure S8**: Western blots of TGFβ1 and integrin $\alpha_V$ expression (biological replicate 3). Top row: total protein, second row: TGFβ1 staining, third row: integrin $\alpha_V$ staining, forth row: sample labels. Integrin $\alpha_V$ staining was done at very low primary antibody dilution, and thus, bands at 120 and 60kDa appeared at the same darkness for many lanes. Only bands at 120kDa were analyzed, as for other replicates.

### S.4 Tissue Pathology

Tissue samples of the heart, brain, kidney, bladder, liver, and spleen were also collected post-euthanasia for histopathological hematoxylin and eosin (H&E) staining to investigate for signs of adverse systemic reactions to either the radiation exposure from CT scanning or the presence of nanoparticles. No signs of pathology were detected due to the devices or nanoparticles. A representative panel of tissues is presented in **Figure S9**.

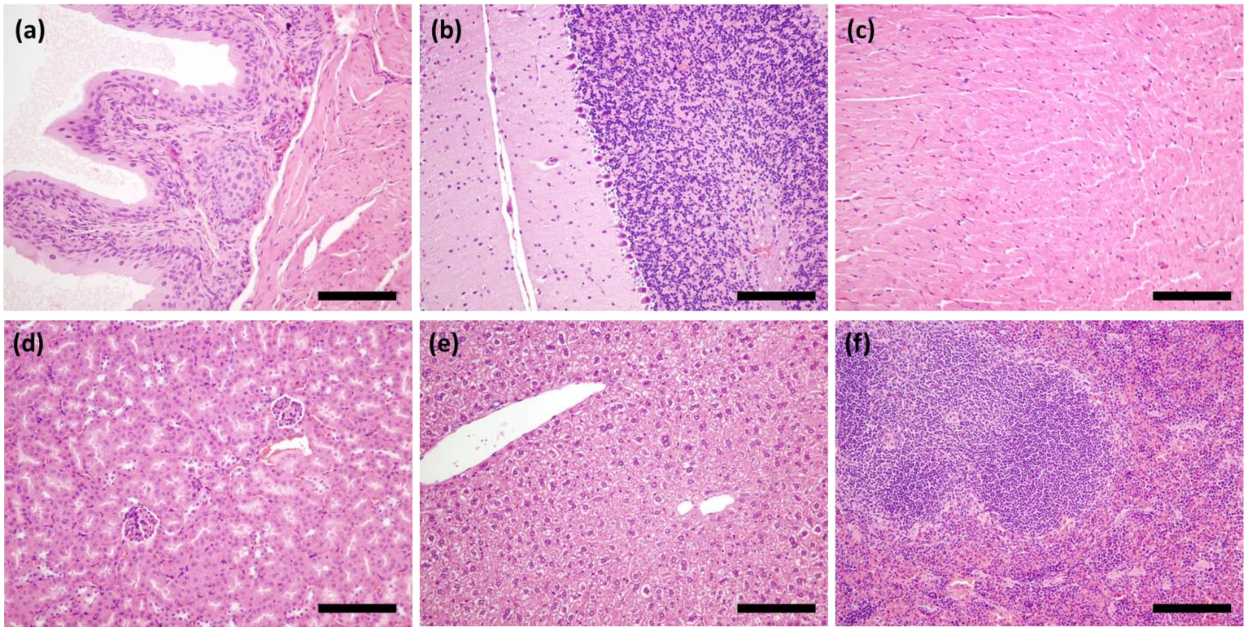


**Figure S9**: Histology of target tissues in the body revealed no abnormalities due to radiation exposure or to the presence of nanoparticles. Example tissues from an animal exposed to serial CT monitoring with an implanted device, demonstrating no abnormalities: (a) bladder, (b) cerebellum, (c) heart, (d) kidney, (e) liver and (f) spleen. Scale bar: 50 µm.

### S.5 In vivo study: Blood Analysis

At the end of the study, following euthanasia, blood was collected from all animals. The blood was used for complete blood cell count (CBC) and clinical pathology. In the CBC analysis, changes were noted in the white blood cell count (WBC) and the relative amount of various blood components, such as monocytes, neutrophils and eosinophils, **Table S1**. The results of clinical pathology are presented in **Table S2**; in some cases, the volume of blood was not sufficient for obtaining a reliable value (marked N/A on the table).

Table S1 Complete blood count of all groups; naïve animals are reference.

| UNIT | Reference | HEMATOLOGY CBC Automated | no device; 0wt% carrageenan; no CT | no device; 1.7wt% carrageenan; no CT | device; 0wt% carrageenan; no CT | device; 1.7wt% carrageenan; no CT | device; 0wt% carrageenan; CT | device; 1.7wt% carrageenan; CT |
|---|---|---|---|---|---|---|---|---|
| g dL$^{-1}$ | 7.3 ± 0.2 | Total Protein (Refractometry) | 7.8 ± 0.1 | 7.0 ± 0.3 | 7.2 ± 0.4 | 7.5 ± 0.25 | 7.7 ± 0.2 | 7.2 ± 0.4 |
| 10x10$^{6}$ µL$^{-1}$ | 10.8 ± 0.1 | RBC | 12.0 ± 0.3 | 10.7 ± 0.2 | 11.1 ± 0.5 | 10.3 ± 0.3 | 11.4 ± 0.4 | 11.1 ± 0.3 |
| g dL$^{-1}$ | 16.9 ± 0.01 | Hgb | 18.9 ± 0.2 | 16.3 ± 0.3 | 17.0 ± 1.4 | 15.7 ± 1.2 | 18.4 ± 0.5 | 17.4 ± 0.6 |
| % | 57.5 ± 0.5 | Hct | 64.0 ± 0.8 | 54.0 ± 1.1 | 56.5 ± 3.8 | 52.6 ± 3.4 | 61.0 ± 2.7 | 58.5 ± 2.1 |
| % | 50.5 ± 0.5 | HCT Spun | 56.3 ± 1.2 | 49.6 ± 1.9 | 50.8 ± 3.8 | 49.4 ± 2.2 | 55.5 ± 3.2 | 54.0 ± 2.1 |
| fL | 53.5 ± 0.5 | MCV | 53.3 ± 0.9 | 50.2 ± 0.7 | 51.0 ± 3.2 | 50.8 ± 1.9 | 53.8 ± 0.8 | 52.5 ± 1.1 |
| pg | 16.0 ± 0.0 | MCH | 15.7 ± 0.5 | 15.0 ± 0.0 | 15.5 ± 1.1 | 15.4 ± 0.8 | 16.0 ± 0.0 | 15.8 ± 0.4 |
| g dL$^{-1}$ | 29.0 ± 0.0 | MCHC | 29.3 ± 0.5 | 30.2 ± 0.4 | 30.2 ± 0.7 | 29.8 ± 0.7 | 30.0 ± 0.8 | 29.8 ± 0.4 |
| g dL$^{-1}$ | 26.5 ± 0.5 | CHCM | 30.0 ± 5.0 | 28.0 ± 0.0 | 27.3 ± 0.5 | 27.0 ± 0.0 | 26.5 ± 0.5 | 27.0 ± 0.6 |
| % | 13.0 ± 0.0 | RDW | 13.7 ± 0.5 | 13.4 ± 0.5 | 14.0 ± 1.4 | 14.0 ± 0.6 | 14.0 ± 0.0 | 14.0 ± 0.0 |
| 10x10$^{3}$ µL$^{-1}$ | 1027.0 ± 30 | Platelet | 562.7 ± 154 | 1018.8 ± 127 | 967.8 ± 221 | 1100.6 ± 183 | 657.0 ± 108 | 928.0 ± 94 |
| fL | 10.3 ± 0.05 | MPV | 10.6 ± 0.2 | 9.0 ± 0.24 | 9.1 ± 0.2 | 9.5 ± 0.2 | 10.2 ± 0.3 | 9.9 ± 0.4 |
| 10x10$^{3}$ µL$^{-1}$ | 8.9 ± 1.3 | WBC | 8.2 ± 1.6 | 9.1 ± 1.8 | 9.5 ± 3.9 | 11.4 ± 2.2 | 7.2 ± 0.3 | 8.0 ± 1.4 |
| **MANUAL** | | **HEMATOLOGY CBC Manual Differential Counts** | | | | | | |
| 10x10$^{3}$ µL$^{-1}$ | 1.5 ± 0.4 | Seg Neut | 1.4 ± 0.2 | 1.6 ± 0.4 | 2.0 ± 1.7 | 2.9 ± 1.6 | 1.3 ± 0.4 | 1.5 ± 0.5 |
| 10x10$^{3}$ µL$^{-1}$ | 0.0 ± 0.0 | Band Neut | 0.0 ± 0.0 | 0.0 ± 0.0 | 0.0 ± 0.1 | 0.0 ± 0.0 | 0.0 ± 0.0 | 0.0 ± 0.04 |
| 10x10$^{3}$ µL$^{-1}$ | 7.3 ± 0.9 | Lymphocyte | 6.4 ± 1.4 | 7.1 ± 1.6 | 6.7 ± 2.1 | 7.1 ± 0.8 | 5.2 ± 0.2 | 6.0 ± 0.7 |
| 10x10$^{3}$ µL$^{-1}$ | 0.2 ± 0.1 | Monocyte | 0.3 ± 0.2 | 0.4 ± 0.12 | 0.5 ± 0.8 | 1.1 ± 0.8 | 0.5 ± 0.1 | 0.5 ± 0.4 |
| 10x10$^{3}$ µL$^{-1}$ | 0.0 ± 0.0 | Eosinophil | 0.2 ± 0.1 | 0.1 ± 0.04 | 0.3 ± 0.2 | 0.3 ± 0.1 | 0.2 ± 0.1 | 0.1 ± 0.1 |
| 10x10$^{3}$ µL$^{-1}$ | 0.0 ± 0.0 | Basophil | 0.0 ± 0.0 | 0.0 ± 0.0 | 0.0 ± 0.0 | 0.0 ± 0.0 | 0.0 ± 0.0 | 0.0 ± 0.0 |
| % | 16.0 ± 2.0 | Seg Neut Pct Manual | 17.0 ± 1.4 | 17.6 ± 3.7 | 19.0 ± 7.7 | 23.8 ± 8.7 | 18.3 ± 4.7 | 18.8 ± 4.7 |
| % | 0.0 ± 0.0 | Band Neut Pct Manual | 0.0 ± 0.0 | 0.0 ± 0.0 | 0.2 ± 0.4 | 0.0 ± 0.0 | 0.0 ± 0.0 | 0.2 ± 0.4 |
| % | 82.0 ± 1.0 | Lymphocyte Pct Manual | 77.0 ± 2.2 | 77.0 ± 5.0 | 73.3 ± 12 | 64.0 ± 8.9 | 73.0 ± 5.5 | 74.8 ± 6.1 |
| % | 2.0 ± 1.0 | Monocyte Pct Manual | 2.7 ± 1.7 | 4.0 ± 1.7 | 4.2 ± 4.5 | 10.0 ± 6.4 | 6.3 ± 1.8 | 5.3 ± 4.6 |
| % | 0.0 ± 0.0 | Eosinophil Pct Manual | 3.3 ± 2.6 | 1.4 ± 0.5 | 3.3 ± 2.0 | 2.2 ± 0.7 | 2.5 ± 1.1 | 0.8 ± 0.9 |
| % | 0.0 ± 0.0 | Basophil Pct Manual | 0.0 ± 0.0 | 0.0 ± 0.0 | 0.0 ± 0.0 | 0.0 ± 0.0 | 0.0 ± 0.0 | 0.0 ± 0.0 |

**Table S2** Clinical pathology of blood for all groups; naïve animals are reference.

| Sample | Urea Nitrogen (mg/dL) | Creatinine (Jaffe) (mg/dL) | Sodium (mmol/L) | Potassium (mmol/L) | Chloride (mmol/L) | TCO2 (mmol/L) | Na/K Ratio | Anion Gap (mmol/L) | Osmolarity Calc (mmol/L) |
|---|---|---|---|---|---|---|---|---|---|
| *naïve* | 13.5 ± 7.5 | <0.1 ± 0.0 | 103.5 ± 54 | 4.9 ± 2.0 | 109.0 ± 0.0 | 11.0 ± 6.0 | 17.5 ± 2.5 | 39.0 ± 0.0 | 219.5 ± 114 |
| *no device; 0wt% carrageenan; no CT* | 20.5 ± 2.1 | <0.1 ± 0.0 | 158.3 ± 3.5 | 9.3 ± 1.3 | 110.3 ± 3.3 | 13.3 ± 2.9 | 17.5 ± 2.2 | 44.0 ± 2.5 | 336.3 ± 8.5 |
| *no device; 1.7wt% carrageenan; no CT* | 21.0 ± 1.2 | 0.2 ± 0.0 | 153.5 ± 2.2 | 8.5 ± 0.6 | 107.3 ± 1.4 | 18.7 ± 1.1 | 18.2 ± 1.6 | 35.8 ± 2.5 | 325.7 ± 3.8 |
| *device; 0wt% carrageenan; no CT* | 19.3 ± 2.6 | 0.2 ± 0.0 | 152.4 ± 1.5 | 8.2 ± 1.0 | 107.8 ± 0.7 | 22.7 ± 3.2 | 19.0 ± 2.4 | 29.3 ± 0.8 | 322.6 ± 3.8 |
| *device; 1.7wt% carrageenan; no CT* | 19.5 ± 2.0 | <0.1 ± 0.0 | 160.3 ± 1.1 | 7.7 ± 0.7 | 110.2 ± 1.5 | 19.0 ± 2.1 | 21.0 ± 1.9 | 38.8 ± 2.3 | 337.2 ± 3.5 |
| *device; 0wt% carrageenan; CT* | 15.8 ± 4.2 | <0.1 ± 0.0 | 141.7 ± 40 | 7.2 ± 2.0 | 114.3 ± 3.7 | 15.4 ± 5.7 | 19.7 ± 1.2 | 40.5 ± 3.6 | 290.0 ± 86 |
| *device; 1.7wt% carrageenan; CT* | 18.5 ± 2.1 | <0.1 ± 0.0 | 151.2 ± 5.2 | 8.6 ± 1.4 | 110.3 ± 2.1 | 16.7 ± 4.8 | 18.0 ± 2.4 | 32.8 ± 7.2 | 319.4 ± 13 |

| Sample | Total Protein (g/dL) | Albumin (g/dL) | Globulin Calc (g/dL) | Glucose (Hexokinase) (mg/dL) | Amylase (U/L) | Total Bili (mg/dL) | Direct Bili (mg/dL) | Indirect Bili (mg/dL) | ALP (U/L) |
|---|---|---|---|---|---|---|---|---|---|
| *naïve* | 4.1 ± 2.1 | 2.3 ± 1.2 | 1.8 ± 0.9 | 140.0 ± 67 | 561.0 ± 316 | 0.2 ± 0.1 | 0.1 ± 0.1 | 0.1 ± 0.0 | 54.0 ± 24 |
| *no device; 0wt% carrageenan; no CT* | 6.2 ± 0.2 | 3.7 ± 0.1 | 2.6 ± 0.2 | 221.3 ± 42 | 759.5 ± 90 | 0.3 ± 0.1 | 0.1 ± 0.1 | 0.2 ± 0.1 | 87.8 ± 5.8 |
| *no device; 1.7wt% carrageenan; no CT* | 6.1 ± 0.6 | 3.2 ± 0.1 | 2.9 ± 0.6 | 201.0 ± 22 | 867.7 ± 290 | 0.2 ± 0.1 | 0.0 ± 0.0 | 0.2 ± 0.1 | 113.2 ± 73 |
| *device; 0wt% carrageenan; no CT* | 6.6 ± 0.4 | 3.0 ± 0.3 | 3.6 ± 0.8 | 190.7 ± 18 | 672.8 ± 67 | 0.2 ± 0.1 | 0.1 ± 0.1 | 0.2 ± 0.1 | 83.2 ± 10 |
| *device; 1.7wt% carrageenan; no CT* | 5.7 ± 0.3 | 3.1 ± 0.1 | 2.6 ± 0.2 | 172.0 ± 36 | 743.8 ± 79 | 0.2 ± 0.0 | 0.0 ± 0.0 | 0.2 ± 0.0 | 78.8 ± 8.2 |
| *device; 0wt% carrageenan; CT* | 5.9 ± 0.4 | 3.4 ± 0.2 | 2.2 ± 0.1 | 124.6 ± 35 | 637.3 ± 158 | 0.4 ± 0.1 | 0.1 ± 0.1 | 0.3 ± 0.1 | 93.2 ± 3.3 |
| *device; 1.7wt% carrageenan; CT* | 5.5 ± 0.2 | 3.2 ± 0.2 | 2.2 ± 0.3 | 213.8 ± 73 | 735.2 ± 63 | 0.4 ± 0.2 | 0.1 ± 0.1 | 0.3 ± 0.1 | 81.3 ± 8.1 |

| Sample | Phosphorus (mg/dL) | Magnesium (mg/dL) | Iron (ug/dL) | AST (U/L) | CK (creatine kinase) (U/L) | Chol (mg/dL) | Calcium (mg/dL) | ALT (U/L) |
|---|---|---|---|---|---|---|---|---|
| *naïve* | 11.2 ± 0.5 | 3.8 ± 0.2 | 205.5 ± 95 | 33.5 ± 16 | 172.5 ± 79 | 117.0 ± 62 | 7.1 ± 3.6 | 20.0 ± 12 |
| *no device; 0wt% carrageenan; no CT* | 11.0 ± 0.5 | 4.2 ± 0.0 | 346.8 ± 10 | 77.3 ± 13 | 397.8 ± 134 | 189.0 ± 17 | 10.5 ± 0.5 | 29.0 ± 3.1 |
| *no device; 1.7wt% carrageenan; no CT* | 10.2 ± 0.6 | 3.5 ± 0.2 | 287.5 ± 13 | 79.3 ± 26 | 372.5 ± 98 | 174.8 ± 28 | 10.6 ± 0.3 | 50.8 ± 36 |
| *device; 0wt% carrageenan; no CT* | N/A | N/A | 290.5 ± 39 | 78.7 ± 40 | 323.5 ± 99 | 157.5 ± 25 | 10.3 ± 0.3 | 32.3 ± 10 |
| *device; 1.7wt% carrageenan; no CT* | 10.5 ± 0.8 | 3.4 ± 0.1 | 285.0 ± 43 | 77.7 ± 33 | 255.7 ± 75 | 148.3 ± 26 | 10.3 ± 0.4 | 50.2 ± 43 |
| *device; 0wt% carrageenan; CT* | N/A | N/A | N/A | 103.2 ± 35 | 340.5 ± 162 | 179.7 ± 23 | 10.5 ± 1.1 | 40.0 ± 15 |
| *device; 1.7wt% carrageenan; CT* | 9.4 ± 0.7 | 3.1 ± 0.3 | 286.3 ± 39 | 60.7 ± 15 | 228.3 ± 84 | 157.2 ± 17 | 10.3 ± 0.8 | 31.2 ± 8.6 |
